# *Synechocystis* KaiC3 displays temperature and KaiB dependent ATPase activity and is important for growth in darkness

**DOI:** 10.1101/700500

**Authors:** Anika Wiegard, Christin Köbler, Katsuaki Oyama, Anja K. Dörrich, Chihiro Azai, Kazuki Terauchi, Annegret Wilde, Ilka M. Axmann

## Abstract

Cyanobacteria form a heterogeneous bacterial group with diverse lifestyles, acclimation strategies and differences in the presence of circadian clock proteins. In *Synechococcus elongatus* PCC 7942, a unique posttranslational KaiABC oscillator drives circadian rhythms. ATPase activity of KaiC correlates with the period of the clock and mediates temperature compensation. *Synechocystis* sp. PCC 6803 expresses additional Kai proteins, of which KaiB3 and KaiC3 proteins were suggested to fine-tune the standard KaiAB1C1 oscillator. In the present study, we therefore characterized the enzymatic activity of KaiC3 as a representative of non-standard KaiC homologs *in vitro*. KaiC3 displayed ATPase activity, which were lower compared to the *Synechococcus elongatus* PCC 7942 KaiC protein. ATP hydrolysis was temperature-dependent. Hence, KaiC3 is missing a defining feature of the model cyanobacterial circadian oscillator. Yeast two-hybrid analysis showed that KaiC3 interacts with KaiB3, KaiC1 and KaiB1. Further, KaiB3 and KaiB1 reduced *in vitro* ATP hydrolysis by KaiC3. Spot assays showed that chemoheterotrophic growth in constant darkness is completely abolished after deletion of Δ*kaiAB1C1* and reduced in the absence of *kaiC3*. We therefore suggest a role for adaptation to darkness for KaiC3 as well as a crosstalk between the KaiC1 and KaiC3 based systems.

**Importance:** The circadian clock influences the cyanobacterial metabolism and deeper understanding of its regulation will be important for metabolic optimizations in context of industrial applications. Due to the heterogeneity of cyanobacteria, characterization of clock systems in organisms apart from the circadian model *Synechococcus elongatus* PCC 7942 is required. *Synechocystis* PCC 6803 represents a major cyanobacterial model organism and harbors phylogenetically diverged homologs of the clock proteins, which are present in various other non-cyanobacterial prokaryotes. By our *in vitro* studies we unravel the interplay of the multiple *Synechocystis* Kai proteins and characterize enzymatic activities of the non-standard clock homolog KaiC3. We show that the deletion of *kaiC3* affects growth in constant darkness suggesting its involvement in the regulation of non-photosynthetic metabolic pathways.

## Introduction

Cyanobacteria have evolved the circadian clock system to sense, anticipate and respond to predictable environmental changes based on the rotation of Earth and the resulting day-night cycle. Circadian rhythms are defined by three criteria: (i) oscillations with a period of about 24 hours without external stimuli, (ii) synchronization of the oscillator with the environment and (iii) compensation of the usual temperature dependence of biochemical reactions, so that the period of the endogenous oscillation does not depend on temperature in a physiological range (2). The cyanobacterial circadian clock system has been studied in much detail in the unicellular model cyanobacterium *Synechococcus elongatus* PCC 7942 (hereafter *S. elongatus*). Its core oscillator is composed of three proteins, which are unique to prokaryotes: KaiA, KaiB, and KaiC (from now on KaiA, KaiB and KaiC; please note that some of the hereafter cited information were gained studying proteins from *Thermosynechococcus elongatus* BP-1 though) (3). The level of KaiC phosphorylation and KaiC’s ATPase activity represent the key features of the biochemical oscillator. KaiA stimulates autophosphorylation and ATPase activity of KaiC, whereas KaiB binding to the complex inhibits KaiA action, stimulates autodephosphorylation activity and reduces ATPase activity of KaiC (4–6). As a consequence of dynamic interactions with KaiA and KaiB, KaiC rhythmically phosphorylates and dephosphorylates with a 24-hour period (3).

KaiC consists of two replicated domains (CI and CII) which assemble into a hexamer forming an N-terminal CI ring and a C-terminal CII ring (7–9). Phosphorylation takes place in the CII ring (10), whereas ATP hydrolysis occurs in both rings (11). In the CII ring, ATP hydrolysis is part of the dephosphorylation mechanism (12, 13). ATP hydrolysis in the CI ring correlates with the period of the clock and temperature compensation and is further required for a conformational change of KaiC, which allows binding of KaiB (5, 14, 15). The levels of phosphorylation and ATP hydrolysis of KaiC serve as the read-out for regulatory proteins, which orchestrate the circadian output (16, 17). For a recent review on the functioning of the KaiABC system, see Swan *et al.* (18).

In a natural day and night cycle, *S. elongatus* orchestrates its metabolism in a precise temporal schedule, with the metabolism being adjusted by the clock but also feeding input information to the clock (19–21). Knockout of the *S. elongatus kai* genes leads to a growth disadvantage. When grown in competition with the wild type under light-dark conditions, the clock deficient mutant cells are eliminated from the culture (22). Growth in light-dark cycles is also impaired by deletion of the regulator of phycobilisome association A (RpaA) (19, 23), the master transcription factor of circadian gene expression. In addition, studies on single nucleotide polymorphisms identified the *rpaA* gene as one of three genes responsible for the faster growth of *Synechococcus elongatus* UTEX 2973 compared to *S. elongatus* (24). Using a transposon library, Welkie *et al.* (25) showed that KaiA, despite being non-essential for growth in light-dark cycles, strongly contributes to the fitness of *S. elongatus* under these conditions. Decreased fitness most likely occurs due to reduced phosphorylation of RpaA in the *kaiA* knockout strain (25). Overall, these data demonstrate the value of the cyanobacterial clock system for metabolic orchestration under natural conditions, with the clock system considered especially important for the transition from light to darkness (26).

Cyanobacteria represent one of the most diverse prokaryotic phyla (27) and little is known about timekeeping mechanisms in other cyanobacteria than *S. elongatus*. A core diurnal genome has been described (28), but temporal coordination varies and, based on genomic analyses, large variations in the cyanobacterial clock systems can be expected (29–34).

The cyanobacterial model strain *Synechocystis* sp. PCC 6803 (from now on *Synechocystis*) contains a standard *kai* operon, encoding homologs of the *kaiA*, *kaiB* and *kaiC* genes (in the following designated *kaiA*_6803_*, kaiB1* and *kaiC1*) as well as two additional copies each of *kaiB* and *kaiC* (designated *kaiB2*, *kaiB3* and *kaiC2, kaiC3* (35). Please note that naming of KaiC3/KaiC2 and KaiB3/KaiB2 is not consistent in the literature. In this paper, we name the *Synechocystis kai* genes according to Aoki and Onai, Wiegard *et al.* and Schmelling *et al.* (28, 30, 32). The *kaiC2* and *kaiB2* genes form an operon, whereas *kaiC3* and *kaiB3* are orphan genes. Based on phylogenetic reconstruction analysis, the KaiB and KaiC proteins of *Synechocystis* were allocated in three phylogenetically different subclasses (32, 36). KaiC1 and KaiB1 display 82 % and 88 % amino acid identity to *S. elongatus* KaiC and KaiB, respectively. Amino acid identity to *S. elongatus* proteins is lower for the other *Synechocystis* Kai homologs, namely, 37 % for KaiC2, 52 % for KaiB2, 55 % for KaiC3 and 48 % for KaiB3. The KaiC homologs show differences in their C-terminus: Both, KaiC2 and KaiC3 show low conservation of the A-loop. KaiC2 further differs from KaiC3 by displaying modified phosphorylation sites (two serine residues instead of serine and threonine in KaiC1 and KaiC3) (32).

Nevertheless, all three KaiC proteins from *Synechocystis* show high conservation of the kinase motif in the CII domain and all three proteins were shown to exhibit autophosphorylation activity (28, 32). KaiA_6803_ stimulates the autophosphorylation of KaiC1, but does not affect the phosphorylation of the two other KaiC homologs. Based on sequence analysis and experimental validation of kinase activity, Schmelling *et al.* (28) concluded that autokinase and autophosphatase activities are highly conserved features of all cyanobacterial KaiC homologs. Likewise, ATPase motifs in the N-terminal CI domain of KaiC are highly conserved in all cyanobacterial KaiC homologs (28). However, ATP hydrolysis was only characterized for several KaiC1 homologs and a KaiC2 homolog from *Legionella pneumophila* (11, 37–39).

In *Synechocystis* deletion of the *kaiAB1C1* gene cluster causes reduced growth in light-dark rhythms, even when grown as single culture, demonstrating a more pronounced phenotype compared to *S. elongatus*, where only a competitive growth disadvantage was observed (40). Deletion of *rpaA* also reduced growth in light-dark cycles, which further implies metabolic orchestration by the *Synechocystis* timer (23). The reported number of oscillating genes in *Synechocystis* varies largely between different studies, which is likely due to differences in growth conditions and strain variations (41–44). Aoki and Onai (30) suggested that KaiC3 and KaiB3 modulate the amplitude and period of the KaiAB1C1 oscillator, whereas disruption of *kaiC2B2* implied a non-clock related function for KaiC2 and KaiB2. There are different *Synechocystis* lab strain variants in use, which show phenotypic variation, for example in their glucose tolerance, motility and tolerance to abiotic stress (45). In the *Synechocystis* strain used in this study (PCC-M, re-sequenced, (46)) the *kaiC2B2* cluster cannot be deleted (40), which further implies a non-clock-related essential function. The function of KaiC3 has not been addressed so far. BLAST analysis identified KaiC3 homologs to be present in addition to KaiC1 in about one third of cyanobacterial genera. KaiC3 homologs were also found in non-cyanobacterial eubacteria and archaea, where they show a higher distribution than the phylogenetically different KaiC1 and KaiC2 homologs (28).

In this study, we therefore aim at a detailed biochemical characterization of the putative clock component KaiC3 and the role of *Synechocystis* KaiB homologs in modulation of KaiC3 function. Characterization of the ATPase activity of KaiC3 was of special interest, because ATP hydrolysis defines the period length and temperature compensation of the Kai oscillator (5). We further demonstrate that the Δ*kaiC3* mutant strain has a growth defect under chemoheterotrophic growth conditions, which is similar, but less pronounced compared to Δ*kaiAB1C1* and Δ*rpaA* strains (23, 40). Our data support the idea of a function of KaiC3 and KaiB3 in fine-tuning the central oscillator composed of KaiA_6803_, KaiB1 and KaiC1 in *Synechocystis*.

(This article was submitted to an online preprint archive (1))

## Results

### Recombinant KaiC3 displays only low ATP synthase activity

Previous bioinformatic analysis predicted that kinase, dephosphorylation and ATPase activities are conserved in KaiC3 (28). So far, only the kinase activity of KaiC3 was experimentally confirmed which differed from the activity of *S. elongatus* KaiC by lacking stimulation by KaiA and KaiA6803 (32). Therefore, we aimed at the characterization of further predicted activities of KaiC3.

The first step in KaiC dephosphorylation is regarded as a reversal of phosphorylation: KaiC transfers its bound phosphoryl group to ADP and thereby synthesizes ATP (12, 13). To investigate reversibility of intrinsic phosphorylation, we tested whether KaiC3 can synthesize [α-^32^P]ATP from [α-^32^P]ADP (Fig. 1). As control, we used phosphorylated and dephosphorylated KaiC (Fig. S1.). Immediately after adding [α-^32^P]ADP, all KaiC proteins started to synthesize [α-^32^P]ATP. Phosphorylated as well as non-phosphorylated KaiCboth showed higher initial ATP synthesis than KaiC3 (Fig. 1). After incubating phosphorylated and dephosphorylated KaiC for 2 hours at 30°C with [α-^32^P]ADP, a relative [α-^32^P]ATP level of 26-28 % was detected. KaiC3 showed low intrinsic ATP synthesis, but formation of relative [α-^32^P]ATP was not significantly higher than in the control with cold ATP. This can be explained by two hypotheses: (i) compared to *S. elongatus* KaiC, KaiC3 showed lower phosphotransferase activity or (ii) higher consumption of the produced ATP. The latter would require that KaiC3 exhibits higher ATPase activity than KaiC. Therefore, we next examined ATP hydrolysis activity of KaiC3.

**Figure 1:**
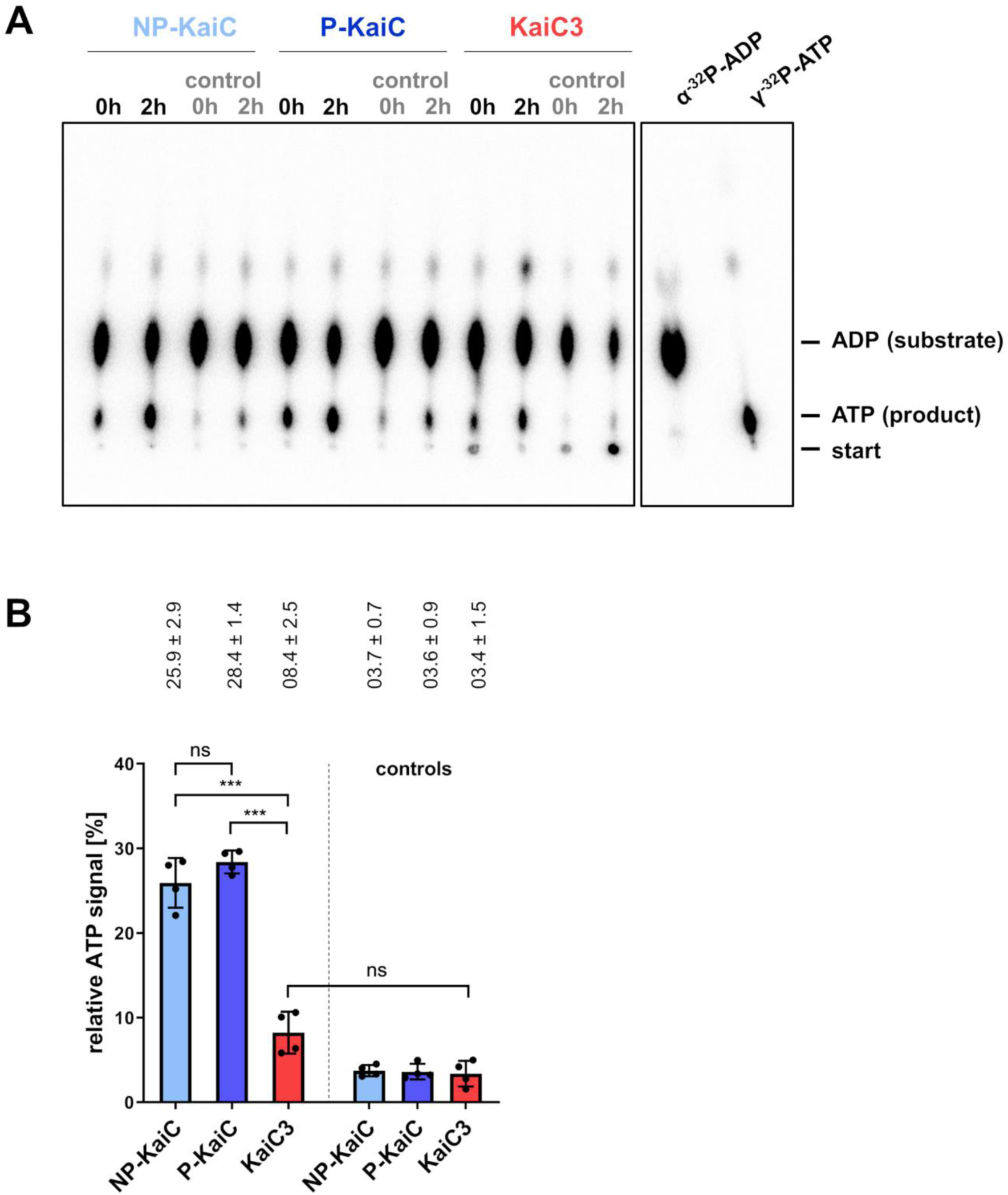
KaiC3 showed no or highly decreased ATP synthase activity compared to KaiC. Prior to the experiment, KaiC was incubated for 2 weeks at 4 °C or overnight at 30 °C to generate fully phosphorylated (P-KaiC) and dephosphorylated (NP-KaiC) protein, respectively (Fig. S1). **A:** Representative autoradiograph of separation of [γ-^32^P]ATP (product) and [α-^32^P]ADP (substrate) via thin layer chromatography after incubation with indicated KaiC proteins for 2h. Controls show the ATP signal in the presence of an excess of cold ADP. For size control [y-^32^P]ATP and [α-^32^P]ADP were separated on the same cellulose F plate. **B:** Relative ATP signals after 2 hours incubation displayed as percentage of all radioactive signals in the corresponding lane (mean with standard deviation from two experiments, each analyzed in duplicates). All values are normalized to the relative ATP signal at 0h incubation time in the presence of an excess of cold ADP. Statistical significance was calculated by Brown-Forsythe and Welch ANOVA tests followed by a Dunnett T3 multiple comparisons test using GraphPad Prism 8. Significance of the mean difference is indicated as follows: non-significant (p ≥ 0.05, ns), significant with p = 0.01 to 0.05 (*), very significant with p = 0.001 to 0.01 (**), extremely significant with p = 0.0001 to 0.001 (***) and extremely significant with p < 0.0001 (****).

### KaiC3 displays ATPase activity

ATP hydrolysis can be measured by quantifying ADP production by KaiC over time (5). Conservation of WalkerA and WalkerB motifs in the CI domain of KaiC3 proteins suggested their capability to hydrolyze ATP (28). We purified recombinant hexameric KaiC3 wild type protein fused to an N-terminal Strep-tag (Strep-KaiC3, Fig. S4) and confirmed the predicted ATPase activity *in vitro*. Based on the measured bulk ATPase activity we calculated that 8.5 ± 1.0 ADP molecules per monomer and day (mean ± s.d., Fig. 2) were produced by Strep-KaiC3. We were not able to purify stable recombinant KaiC1 with sufficient purity and therefore used the highly similar KaiC ortholog from *S. elongatus* for comparison. The ATPase activity of Strep-KaiC3 was about 45 % of the value for KaiC from *S. elongatus* (19.1 ± 3.3 ADP per day (5)). Hence, the above-observed lower level of net ATP production by KaiC3 was not based on a higher ATP consumption, but due to lower dephosphorylation activity *per se*. In KaiC, the ATPase activity changes after substitution of the phosphorylation sites (5). We therefore generated variants of Strep-KaiC3, in which residues S423 and T424 are either replaced with aspartate and glutamate (Strep-KaiC3-DE) or replaced with two alanine residues (Strep-KaiC3-AA). Strep-KaiC3-AA, showed more than two fold increased ATPase activity, whereas ATP hydrolysis by Strep-KaiC3-DE was only slightly different from the wild-type protein (Fig. 2) and not reduced as reported for *S. elongatus* KaiC-DE (5).

**Figure 2:**
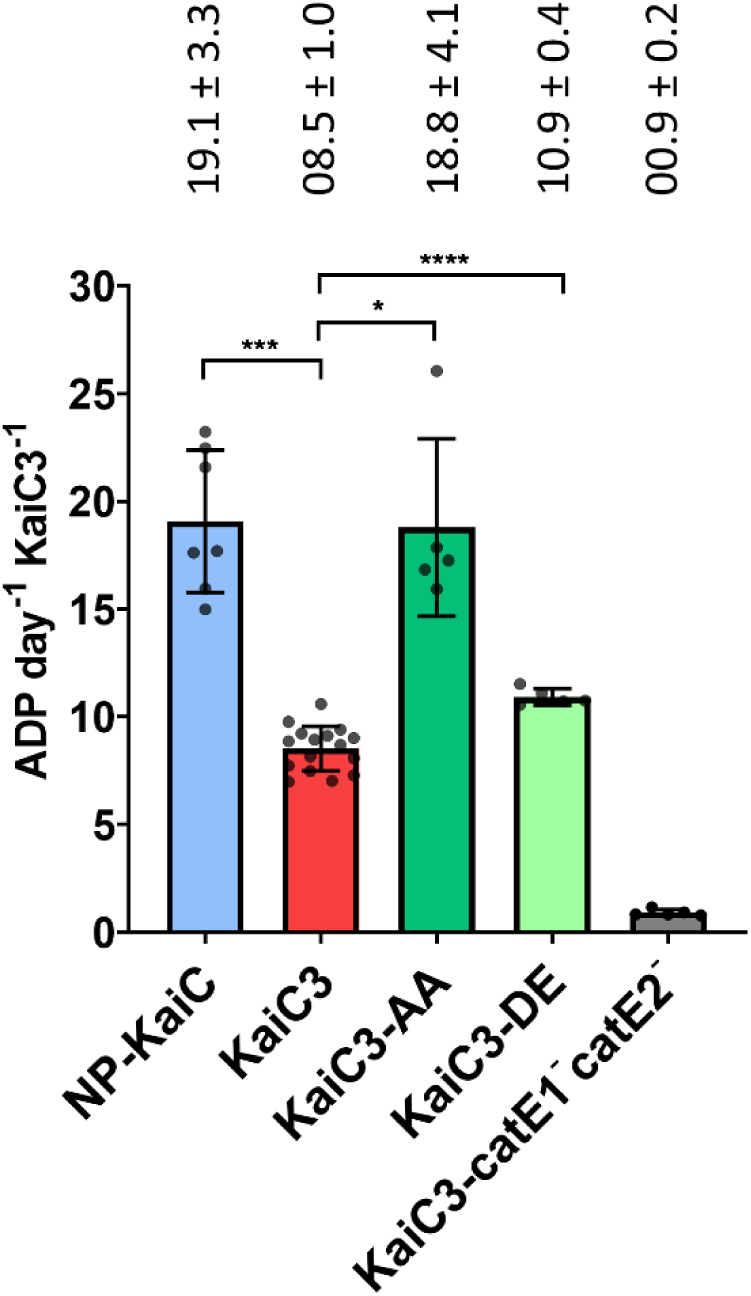
ATPase activity of Strep-KaiC3 and variants Strep-KaiC3-DE and Strep-KaiC3-AA. Strep-KaiC3, Strep-KaiC3-DE and Strep-KaiC3-AA were incubated for 24 hours at 30 °C and ADP production per day and per Strep-KaiC3 monomer was calculated. As a control we used Strep-KaiC3-catE1^−^catE2^−^, which lacks both ATPase motifs due to replacement of catalytic glutamates 67,68, 310 and 311 with glutamine residues. For comparison ADP production by KaiC was monitored for 24 or 48 hours. Strep-KaiC was dephosphorylated prior to the experiment by incubation at 30°C overnight. Shown are mean values with standard deviation of at least five replicates. Statistical significance was calculated by Brown-Forsythe and Welch ANOVA tests followed by a Dunnett T3 multiple comparisons test using GraphPad Prism 8. Significance of the mean difference is indicated as follows: non-significant (p ≥ 0.05, ns), significant with p = 0.01 to 0.05 (*), very significant with p = 0.001 to 0.01 (**), extremely significant with p = 0.0001 to 0.001 (***) and extremely significant with p < 0.0001 (****). Please note that we averaged all measurements for Strep-KaiC3 WT at 30 °C and therefore repeatedly display them in Fig. 2, Fig. 4A and Fig. 4B.

### KaiC3 interacts with KaiB3 and components of the standard KaiAB1C1 oscillator

*In vitro* and *in silico* studies suggested that KaiA_6803_, KaiB1 and KaiC1 form the standard clock system of *Synechocystis* (30, 32). Since Aoki and Onai (30) suggested that KaiC3 might modulate the main oscillator function, we performed protein-protein interaction studies in order to reveal a possible crosstalk between the multiple Kai proteins. First, interaction was determined by yeast-two hybrid analysis using KaiC1, KaiC3, KaiB1 and KaiB3 fused to AD and BD domains, respectively. The color change of the colonies indicates β-galactosidase activity and is an estimate for the interaction of the respective two proteins. As expected, these experiments showed self-interaction of KaiC3 (Fig. 3A), since KaiC homologs are known to form hexamers (7, 9). In addition, an interaction of KaiC3 with KaiB3 was detected (Fig. 3A) which is in line with bioinformatic analysis showing frequent co-occurrence of *kaiC3* and *kaiB3* genes in genomes (28). We could not detect an interaction between KaiC3 and KaiA_6803_ by yeast-two hybrid analysis (Fig. 3B), but detected a heteromeric interaction between KaiC1 and KaiC3 using different protein fusion variants (Fig. 3C). These results confirm previously published data, which showed (i) co-purification of KaiA with KaiC1 but not with KaiC3 and (ii) a weak interaction between the two KaiC homologs KaiC1 and KaiC3 in ex-vivo pulldown analysis followed by Western Blot analysis (32). Besides KaiC1, also the cognate KaiB protein, KaiB1, showed an interaction with KaiC3 in our analysis (Fig. 3C), corroborating the hypothesis that there is a cross talk between the putative KaiC3-B3 system and the core oscillator KaiAB1C1. Such a possible cross talk via the KaiB proteins was further supported by our *in vitro* pull-down assays, in which KaiC3 interacted with KaiB3 (Fig. S3B,D) and further with KaiB1 (Fig. S3B,D). Also KaiC1 interacted with both, KaiB1 and KaiB3 homologs (Fig. S3,A,C). One must take into account that based on the *in vitro* pull-down assays using *Synechocystis* whole cell extracts we cannot exclude indirect interactions. As we have shown that KaiC1 and KaiC3 interact with each other, both proteins could bind as a hetero-hexamer to the GST-tagged KaiB proteins and therefore co-elute from the affinity matrix.

**Figure 3:**
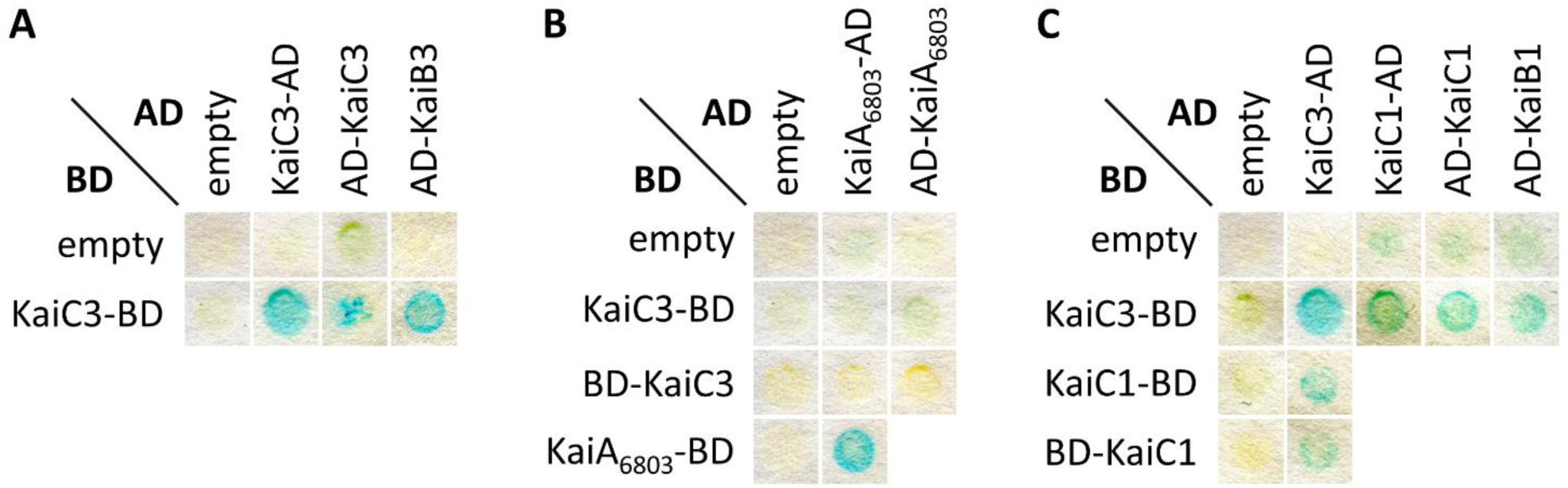
KaiC3 interacts with KaiB3 and the proteins of the main oscillator KaiC1, KaiB1. **A, B, C:** Yeast two-hybrid reporter strains carrying the respective bait and prey plasmids were selected by plating on complete supplement medium (CSM) lacking leucine and tryptophan (-Leu -Trp). Physical interaction between bait and prey fusion proteins is indicated by a color change in the assays using 5-brom-4-chlor-3-indoxyl-β-D-galactopyranoside. **AD**: GAL4 activation domain; BD, GAL4 DNA-binding domain. Shown are representative results of two replicates. For clear presentation, spots were assembled from several assays (original scans are shown in Fig. S2).

### ATPase activity of KaiC3 is reduced in the presence of KaiB1 and KaiB3

The interaction of KaiC3 with KaiB1 and KaiB3 (Fig. 3 and Fig. S3) suggested a regulation of KaiC3 activity by these two KaiB proteins. We therefore measured ATP hydrolysis by Strep-KaiC3 in the presence of 0.04 mg ml^−1^ KaiB1 and KaiB3 proteins, respectively (Fig. 4A). After size exclusion chromatography KaiB1 was only eluted as a tetramer (44 and 72 kDa, respectively, depending on the column), whereas KaiB3 was eluted as monomer (13/23 kDa) and tetramer (41/70 kDa) (Fig. S4). Therefore, the monomeric and tetrameric KaiB3 fractions were tested separately. In all measurements, the ATPase activity was linear and showed no oscillations. ATP hydrolysis was reduced by 55 % after the addition of the KaiB3 monomer, but not affected by the KaiB3 tetramer. The KaiB1 tetramer also inhibited the ATPase activity of Strep-KaiC3 (35 % reduction compared to Strep-KaiC3 alone, Fig. 4). These data imply a trend, that KaiB1 has less effect on KaiC3’s ATPase activity than the KaiB3 monomer, but the differences were not statistically significant (Fig. 4). Because ATPase activity of KaiC is influenced by KaiA (5), we also investigated the effect of KaiA_6803_ on KaiC3, but in line with the lack of interaction in our yeast two-hybrid analysis data (Fig. 3B), ATPase activity of Strep-KaiC3 was not significantly affected by KaiA_6803_ (Fig. 4A). Fig. 4C summarizes the Kai protein interactions and regulation of KaiC3’s ATPase activity by KaiB proteins.

**Figure 4:**
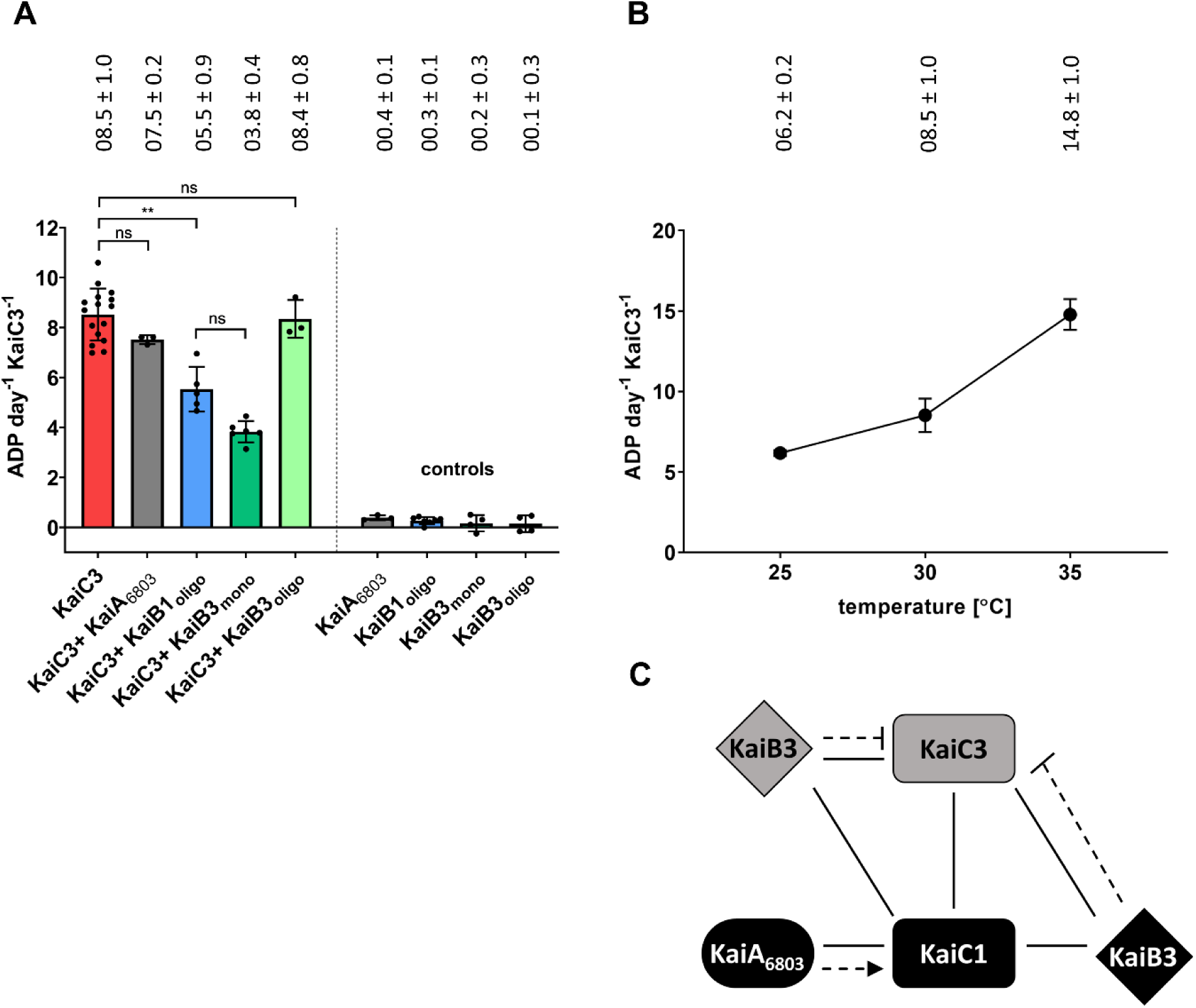
ATPase activity of Strep-KaiC3 is regulated by KaiB proteins and is temperature dependent. **A:** ATPase activity of Strep-KaiC3 was decreased in the presence of KaiB1 and monomeric KaiB3 but was not influenced by oligomeric KaiB3 and by KaiA_6803_. Shown are mean values with standard deviation of at least three replicates. Controls show that KaiA_6803_ and KaiB proteins alone did not display ATPase activity (referring to Strep-KaiC3 monomer activity). Statistical significance was calculated by Brown-Forsythe and Welch ANOVA tests followed by a Dunnett T3 multiple comparisons test using GraphPad Prism 8. Significance of the mean difference is indicated as follows: non-significant (p ≥ 0.05, ns), significant with p = 0.01 to 0.05 (*), very significant with p = 0.001 to 0.01 (**), extremely significant with p = 0.0001 to 0.001 (***) and extremely significant with p < 0.0001 (****). **B:** ATPase activity of Strep-KaiC3 is temperature-dependent. Strep-KaiC3 was incubated for 24 hours at the indicated temperatures and ADP production per day and monomer Strep-KaiC3 was calculated. Shown are mean values with standard deviation of at least three replicates. A temperature-compensated activity would result in flat horizontal line. Please note that we averaged all measurements for Strep-KaiC3 WT at 30°C and therefore repeatedly display them in Fig. 2, Fig. 4A and Fig. 4B. **C:** Summary of the crosstalk between the KaiAB1C1 standard oscillator and KaiC3 in *Synechocystis*. We hypothesize that KaiC1 forms a standard oscillator together with KaiA_6803_ and KaiB1. KaiC3 might form an additional regulatory mechanism together with KaiB3. The two systems intertwine by interactions of the KaiC proteins with each other and the non-corresponding KaiB homologs. Solid lines indicate experimentally verified physical interactions (this paper and Wiegard *et al*.(32)), dotted arrows indicate stimulation of phosphorylation (32) and dotted lines with bars show inhibition of ATPase activity.

### ATPase activity of KaiC3 is temperature dependent

True circadian clocks are characterized by temperature compensated oscillations, which ensure robust time measurements under temperature fluctuations. In the *S. elongatus* KaiABC clock, overall temperature compensation is derived from KaiC’s ATPase activity, which is stable between 25 and 35 °C (Q_10_=1.2, (5)). We therefore asked whether N-Strep-KaiC3, as a representative of non-standard KaiC homologs, shows temperature compensation as well. Measurements at 25, 30 and 35 °C, however, revealed a temperature-dependent ADP production by KaiC3 (Q_10_=2.4, Fig. 4B; activation energy = 66.5 kJ mol^−1^, Fig. S5A). Hence, KaiC3 is lacking a characteristic feature of circadian oscillators. In accordance, dephosphorylation of Strep-KaiC3 was higher at 25 °C than at 30 and 35 °C (Fig. S5B)

### The non-standard KaiC3 protein supports growth of *Synechocystis* cells in the dark

As *in vitro* analyses suggested an interaction of KaiC3 with KaiB1 and KaiC1, we aimed at elucidating a putative role of KaiC3 and its crosstalk with the KaiC1-based system in the cell. Clock factors are reported to be essential for cell viability in light-dark cycles (19, 23, 25). For assessing growth of mutant strains lacking *kai* genes, we used a spot plating assay (48). In this assay, the amount of cells plated on agar in serial dilutions is compared to the amount of cells which are able to form a colony. The plating efficiency then allows us to compare the sensitivity of cells to changing light conditions. Notably, this method allows only limited assertions on the growth rate of different strains. Deletion of the *kaiAB1C1* cluster of *Synechocystis* resulted in lower cell viability in light-dark cycles (40). This was even more pronounced under photomixotrophic (0.2 % glucose) compared to photoautotrophic conditions, whereas deletion of *kaiC3* had no effect on cell viability in light-dark cycles (40). As KaiC3 homologs are also present in non-photosynthetic bacteria (28), we were interested in the growth of the Δ*kaiC3* strain in the dark. The *Synechocystis* wild-type strain used here, in contrast to previous studies (47), is able to grow in complete darkness when supplemented with glucose (23). We therefore analyzed the viability of Δ*kaiC3* cells via spot assays in constant light and in complete darkness on agar plates containing 0.2 % glucose (Fig. 5A). Under photomixotrophic growth conditions with continuous illumination, wild type and Δ*kaiC3* showed similar viability, whereas the viability of the Δ*kaiAB1C1* strain was reduced, reflected by a lower plating efficiency. Further, in complete darkness, wild type displayed detectable growth, while growth of Δ*kaiAB1C1* was abolished completely. Sensitivity of the Δ*kaiAB1C1* mutant to mixotrophic conditions in the light and lack of growth in the dark highlights the importance of the KaiAB1C1 oscillator for the switch between these two different metabolic modes of *Synechocystis*. The Δ*kaiC3* strain could grow in the dark but showed some impairments. The plating efficiency in serial dilutions indicates similar viability of the wild-type and Δ*kaiC3* deletion strain (Fig. 5A). However, dark growth seemed to be more impaired in the Δ*kaiC3* strain when compared to the wild type. For a more quantitative assessment, intensity measurements of single spots of the *Synechocystis* wild type and *kaiC3* deletion strain were performed (Fig. 5B), validating the growth impairment of the *kaiC3* deletion strain in complete darkness. Thus, KaiC3 seems to be linked to dark adaption of *Synechocystis* cells, yet not as essential as the core oscillator KaiAB1C1.

**Figure 5:**
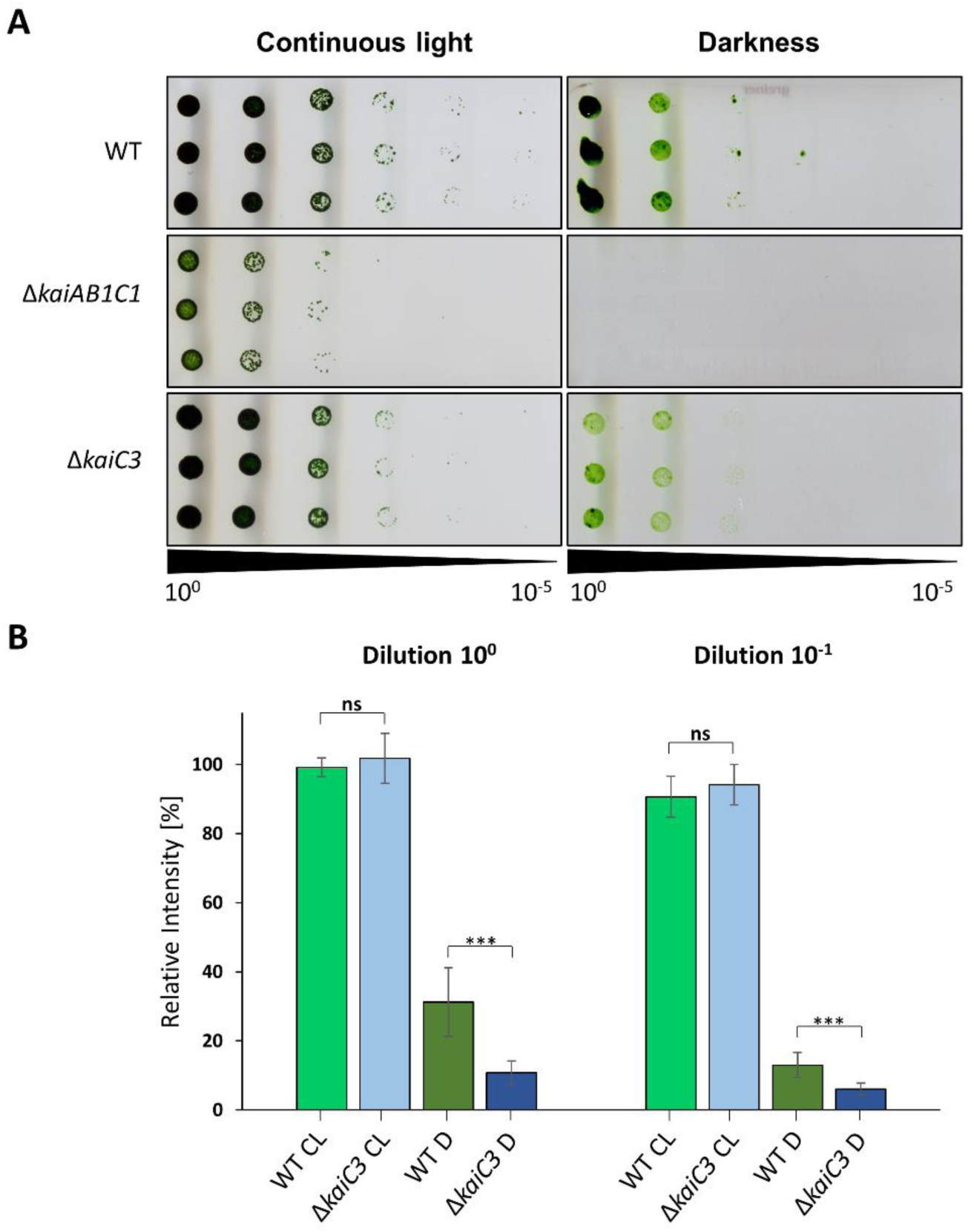
The Δ*kaiC3* strain shows growth defects in complete darkness. **A:** Proliferation of the *Synechocystis* wild type, the *kaiAB1C1* deletion mutant (Δ*kaiAB1C1*) and the *kaiC3* deletion mutant (Δ*kaiC3*) was tested under mixotrophic conditions in continuous light and under heterotrophic dark conditions. Strains were grown in liquid culture under constant light and different dilutions were spotted on agar plates containing 0.2 % glucose. Plates were analyzed after further incubation for 6 or 28 days of continuous light and darkness, respectively. A representative result of three independent experiments is shown. **B:** Measurement of single spot densities of the *Synechocystis* wild-type and *kaiC3* deletion strains. Single spot intensities were determined with Quantity One (Bio-Rad) and normalized to the intensity of the wild-type spot of the respective control (continuous light condition). Shown are mean values with standard deviation of at least three replicates. Statistical significance was calculated by Brown-Forsythe and Welch ANOVA tests followed by a Dunnett T3 multiple comparisons test using GraphPad Prism 8. Significance of the mean difference is indicated as follows: non-significant (p ≥ 0.05, ns), significant with p = 0.01 to 0.05 (*), very significant with p = 0.001 to 0.01 (**), extremely significant with p = 0.0001 to 0.001 (***) and extremely significant with p < 0.0001 (****). CL, continuous light; D, darkness.

## Discussion

### Characterization of enzymatic activities of KaiC3

Previous sequence analysis suggested that the three enzymatic activities of KaiC, which are autophosphorylation, autodephosphorylation and ATPase activity, are conserved in all cyanobacterial KaiC proteins (28). This hypothesis is supported by experiments demonstrating autophosphorylation of all *Synechocystis* KaiC proteins (32), different cyanobacterial KaiC1 and KaiC3 homologs (28, 37, 39), and even non-cyanobacterial KaiC homologs from *Rhodopseudomonas palustris* (38), *Legionella pneumophila* (48) as well as two thermophilic Archaea (28). In the current paper, we provide experimental evidence that ATPase activity is conserved in KaiC3 as well. However, the level of the ATPase activity seems to vary between KaiC proteins. While ATPase activity of the KaiC from *Prochlorococcus marinus* MED4 was reported to have the same activity as KaiC (37), we show here that the ATPase activity of KaiC3 is lower. In contrast, ATP hydrolysis activity of the KaiC2 homolog from *Legionella pneumophila* and KaiC1 homolog from *Gloeocapsa* sp. PCC 7428 were reported to be elevated when compared to the standard KaiC protein (38, 39).

ATPase activity of KaiC3 further differed from that of KaiC, by lacking temperature compensation (Fig. 4). Therefore, KaiC3 cannot be the core component of a true circadian oscillator. This is further supported by differences in the ATP hydrolysis rate between the phosphomimetic variants of KaiC and KaiC3. ATPase activity of the true clock protein KaiC from *S. elongatus* depends on its phosphorylation status. In the KaiC-AA variant, which mimics the dephosphorylated state of KaiC, the ATPase activity is higher than average, whereas, in KaiC-DE, mimicking the phosphorylation state, it is diminished (5). In our analysis, ATP hydrolysis by Strep-KaiC3-AA was similarly increased in comparison to Strep-KaiC3. However, Strep-KaiC3-DE did not show reduced activity as was shown for KaiC-DE, implying that the phosphorylation state of KaiC3 does not influence KaiC3’s ATPase activity.

The ATPase activity of Strep-KaiC3 was affected decisively by the addition of KaiB3 and KaiB1. This points at a putative mechanism how KaiC3 might fine-tune the standard oscillator: Binding of KaiB1 to KaiC3 will reduce the free KaiB1 concentration in the cell and thereby influence the KaiAB1C1 oscillator. Both KaiB homologs led to decreased ATPase activity of Strep-KaiC3 (Fig. 4A). One needs to take into consideration that we used 0.04 mg ml^−1^ KaiB protein for all assays. As KaiB1 was present only as tetramer in the used fractions, this resulted in a molar concentration of 0.82 µM KaiB1 tetramers versus 3.36 µM KaiB3 monomers, respectively. *S. elongatus* KaiB and KaiB3 both exist as monomer or tetramer in solution and KaiB is known to bind as a monomer to KaiC, (49–51). In our analysis where we separated monomeric and tetrameric KaiB3 forms, only monomeric KaiB3 showed an effect on ATP hydrolysis by KaiC3. Therefore, addition of a similar monomeric concentration (3.28 µM KaiB1 and 3.36 µM KaiB3) was important to compare the effect of KaiB1 and KaiB3. As a tetramer, KaiB subunits adopt a different unique fold, which was also observed in crystals of KaiB1 (52). Residues K57, G88 and D90, which are important for fold switching and hence transition between tetrameric and monomeric KaiB (51), are also conserved in KaiB3. We, therefore, cannot exclude that addition of a higher molar concentration of KaiB3 tetramers might affect ATPase activity of KaiC3. However, we show only *in vitro* data here. In the cell, protein concentrations as well as spatial and temporal separation of the different proteins might have an influence on these interactions. In addition, we do not take possible heteromeric interactions among the KaiB proteins into account.

### Function of KaiC3 in an extended network

The interaction studies shown here imply a crosstalk between KaiC1 and KaiC3 (Fig. 4C). KaiC1 is believed to form a standard oscillator together with KaiA_6803_ and KaiB1 (30, 32), which is supported by the KaiC1-KaiB1 interaction observed here (Fig. 3C, Fig. S3). We further hypothesize that KaiC3 acts in a separate non-circadian regulatory system together with KaiB3. In accordance with bioinformatic analysis, which showed a significant co-occurrence of KaiB3 and KaiC3 in Cyanobacteria (28), KaiB3 had a stronger effect on the ATPase activity of KaiC3 than KaiB1 (Fig. 4A). In the cell, the KaiAB1C1 and KaiB3C3 systems could work independently from each other. However, KaiC3 is able to form hetero-oligomers with KaiC1 and was shown to interact with KaiB1 as well. Therefore, interference between the KaiC3-KaiB3 system and the proteins of the standard oscillator is very likely. Further, we exclude that KaiA_6803_ is involved in the putative crosstalk by following reasons: (i) KaiA_6803_ did neither stimulate ATP hydrolysis nor kinase activity of KaiC3 (this paper and Wiegard *et al.* (32)); (ii) We were not able to show an interaction of KaiC3 and KaiA_6803_ using different approaches (this paper and Wiegard *et al.* (32)); (iii) KaiA interacting residues are not conserved in cyanobacterial KaiC3 homologs, and (iv) KaiC3 homologs are present in organisms which do not harbor KaiA (28).

The growth defect of the mutant strains, Δ*kaiAB1C1* and Δ*kaiC3,* in complete darkness suggests that the here proposed putative KaiC3-KaiB3 system and the KaiAB1C1 oscillator may target similar cellular functions. The metabolic network of cyanobacteria is described as temporally partitioned with extensive effects of day-night transitions, involving shifts in ATP and reductant levels and alterations of the carbon flux (25). In *S. elongatus*, environmental signals can be fed into the main clock output system via the transcriptional regulator RpaB (53). In contrast to *S. elongatus*, *Synechocystis* is able to grow chemoheterotrophically, which adds another layer of complexity to day-night transitions and demand for further regulatory elements. The observed impaired growth of the *kaiC3* mutant in darkness supports the idea of KaiC3 and KaiB3 functioning as such additional elements to adjust the state of the main *Synechocystis* oscillator. Conversely, it is also possible that KaiC3 function is controlled by the KaiAB1C1 clock system. Köbler *et al.* (23) demonstrated that solely KaiC1, but not KaiC3, interacts with the main output histidine kinase SasA in the *Synechocystis* timing system. Thus, in *Synechocystis*, only the main oscillator feeds timing information into the SasA-RpaA output system to control the expression of many genes involved in dark growth (23). The output pathway for KaiC3 is unknown so far and it might be possible that the only function of KaiC3 is to modulate the function of the main oscillator in response to a yet unknown input factor.

## Material and Methods

### Cloning, expression and purification of recombinant Kai proteins

Genes encoding KaiB1 (ORF *slr0757*) and KaiB3 (ORF *sll0486*) were amplified from genomic *Synechocystis* wild type DNA using specific primers (Table S1) and Phusion Polymerase (New England Biolabs). After restriction digest with BamHI and NotI, amplified fragments were inserted into pGEX-6P1 (GE Healthcare) and the resulting plasmids were used for heterologous expression. For production of recombinant KaiC as well as KaiA_6803_, KaiB1, KaiB3 and KaiC3, we used pGEX-based plasmids described in Wiegard *et al.* (32) (see also Table S2 for a list of all plasmids used in this study). A detailed protocol of expression and purification can be found on protocols.io (https://dx.doi.org/10.17504/protocols.io.48ggztw). Briefly, proteins were expressed as GST-fusion proteins in *E. coli* BL21 [DE3] or NEB Express (New England Biolabs) and lysed in 50 mM Tris/HCl (pH8), 150 mM NaCl, 0.5 mM EDTA, 1 mM DTT (+ 5 mM MgCl_2_, 1 mM ATP for KaiC proteins). Purification was performed via *batch* affinity chromatography using glutathione agarose 4B (Macherey and Nagel) or glutathione sepharose 4B (GE Healthcare) in the same buffer. Finally, the GST-tag was cleaved off using PreScission protease (GE Healthcare) in 50 mM Tris/HCl (pH8), 150 mM NaCl, 1 mM EDTA, 1 mM DTT (+ 5 mM MgCl_2_, 1 mM ATP for KaiC proteins). If homogeneity of the proteins was not sufficient, they were further purified via anion exchange chromatography using a MonoQ or ResourceQ column (GE Healthcare Life Sciences) using 50 mM Tris/HCl (pH8), 1 mM EDTA, 1 mM DTT (+ 5 mM MgCl_2_, 1 mM ATP for KaiC proteins) and a 0-1M NaCl gradient.

To produce Strep-KaiC3 variants with amino acid substitutions, the *kaiC3* gene in the pGEX-kaiC3 vector was modified by site directed mutagenesis using the Quick-Change Site-Directed Mutagenesis Kit (Stratagene) or Q5 Site-Directed Mutagenesis Kit (New England Biolabs). Base triplets encoding S423 and T424 were changed to code for alanine or for aspartate and glutamate resulting in *kaiC3-AA* and *kaiC3-DE* genes, respectively. To generate *kaiC3-catE1^−^catE2*, two subsequent site-directed mutagenesis reactions were performed to exchange nucleotides encoding E67 and E68 as well as nucleotides encoding E310 and E311 all with bases encoding glutamine (all primers used for mutagenesis are listed in Table S1). Afterwards, *kaiC3* WT and modified *kaiC3* genes were amplified with KOD-Plus-Neo polymerase (Toyobo) using pASK-kaiC3 primers (Table S1). Amplicons were digested with SacII and HindIII and ligated into the respective restriction sites of pASK-IBA5plus (IBA Life sciences). For purification of recombinant Strep-KaiC, the pASK-IBA-5plus based vector described in Oyama *et al.* (15) was used. Strep-KaiC3 proteins were expressed in *E. coli* Rosetta gamiB (DE3) or Rosetta gami2 (DE3) cells (Novagen). Expression of Strep-KaiC was carried out in *E. coli* DH5α. Cells were cultured in LB medium containing 100 μg ml^−1^ ampicillin with vigorous agitation at 37 °C. At OD_600nm_ 0.33-0.68 protein expression was induced with 200 ng ml^−1^ anhydrotetracycline and the strains were further incubated as following: Strep-KaiC: 7h at 37°C, Strep-KaiC3-WT: 5h at 35°C or 3.5h at 37°C, Strep-KaiC3-AA and Strep-KaiC3-catE1^−^catE2^−^: 18°C overnight, Strep-KaiC3-DE: 25°C overnight. Cells were harvested and lysed by sonication in ice-cold buffer W [20mM Tris/HCl (pH8), 150 mM NaCl, 5 mM MgCl_2_, 1 mM ATP (+ 2 mM DTT for Strep-KaiC3 proteins)] including protease inhibitors (protease inhibitor cocktail, Roche or Nacalai). Soluble proteins were loaded on self-prepared columns packed with Strep-Tactin XT superflow or Strep-Tactin Sepharose (IBA lifesciences) and purified under gravity flow. After washing with buffer W, Strep-KaiC proteins were eluted with ice cold buffer W + 50 mM D(+)biotin (for Strep-Tactin XT Superflow) or ice cold buffer W + 2.5 mM desthiobiotin (for Strep tactin Superflow). See https://dx.doi.org/10.17504/protocols.io.meac3ae for a detailed protocol.

All proteins used for ATPase activity measurements were further purified via size exclusion chromatography. Strep-KaiC3 and Strep-KaiC proteins were applied on a Sephacryl S300 HR HiPrep 16/60 Sephacryl column (GE Healthcare) and separated in 20 or 50 mM Tris/HCl (pH 8.0), 150 mM NaCl, 2 mM DTT, 1 mM ATP and 5 mM MgCl2. For separation of KaiB1 and KaiB3, a Sephacryl S200 HR HiPrep 16/60 column (GE Healthcare) or Superdex 200 Increase 10/30 GL column (GE Healthcare) and 20 or 50 mM Tris/HCl (pH 8.0), 150 mM NaCl, 2 mM DTT as running buffer were used. KaiA_6803_ was purified on a Superdex 200 Increase 10/30 GL column (GE Healthcare) in 20 mM Tris/HCl (pH 8.0), 150 mM NaCl, 2 mM DTT. See https://dx.doi.org/10.17504/protocols.io.mdtc26n for further details.

### ATPase activity

For ATPase measurements, KaiC proteins fused to an N-terminal Strep-tag were used and all Kai proteins were purified via size exclusion chromatography (see above). 3.45 μM of Strep-KaiC3 variants were incubated in ATPase buffer (20 mM Tris/HCl (pH8), 150 mM NaCl, 1 mM ATP, 5 mM MgCl_2_) at 25 °C, 30 °C or 35 °C for 24 hours. To analyze the influence of KaiA_6803_ and KaiB proteins on KaiC3 ATPase activity, we mixed 0.2 mg ml^−1^ KaiC3 with 0.04 mg ml^−1^ KaiA_6803_ or 0.04 mg ml^−1^ KaiB and incubated the mixtures for 24 hours at 30 °C. Monomeric and oligomeric KaiB3 were analyzed separately. To monitor ADP production, every 3, 4 or 6 hours 2 μl of the reaction mixture were applied on a Shim-Pack-VP-ODS column (Shimadzu) and separated using 100 mM phosphoric acid, 150 mM triethylamine, 1 % acetonitrile as running buffer. ADP production per monomer KaiC and 24 hours was calculated using a calibration curve. A detailed protocol can be found on protocols.io (https://dx.doi.org/10.17504/protocols.io.mebc3an). The Q_10_ value was calculated from ATPase measurements at 25 °C and 35 °C, using the formula 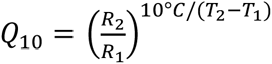

### ATP synthase activity

To investigate dephosphorylation via ATP synthesis, we used KaiC proteins which were expressed as GST-fusion proteins and subsequently cleaved off their GST-tag. 3 μM KaiC in ATP synthesis buffer (20 mM Tris/HCl(pH8), 150 mM NaCl, 0.5 mM EDTA, 5 mM MgCl_2_, 0.5 mM ATP) was mixed with 0.8 µCi ml^−1^ [α^32^P]ADP and stored at −20 °C or incubated for 2 hours at 30 °C. As a control, the same experiment was performed in the presence of 0.5 mM ADP. After 20-fold dilution, nucleotides in a 0.5 μl reaction mixture were separated via thin layer chromatography using TLC PEI Cellulose F plates (Merck Millipore) and 1 M LiCl as solvent. [α-^32^P]ADP and [γ-^32^P]ATP were separated in parallel to identify the signals corresponding to ADP and ATP, respectively. Dried plates were subjected to autoradiography and signals were analyzed using a Personal Molecular Imager FX system (Bio-Rad) and ImageLab software (Bio-Rad). For each reaction mixture, the relative intensity of [α-^32^P]ATP was calculated as percentage of all signals in the corresponding lane. Because [α-^32^P]ATP was already synthesized during mixing of the samples, the relative ATP intensity measured in the −20 °C sample containing 0.5 mM ADP was subtracted for normalization. The principle of this method is based on Egli *et al.* (13). A detailed protocol is available on protocols.io (https://dx.doi.org/10.17504/protocols.io.48qgzvw).

### Yeast two-hybrid assays

For yeast two-hybrid assays, vectors containing the GAL4 activation domain (AD) and the GAL4 DNA-binding domain (BD) were used. Genes of interest were amplified from *Synechocystis* wild type genomic DNA with the Phusion Polymerase (NEB) according to manufacturer’s guidelines. Indicated restriction sites were introduced via the oligonucleotides listed in Table S1. Vectors and PCR fragments were cut with the respective restriction enzymes (Thermo Fisher Scientific) and the gene of interest was ligated into the vector, leading to a fusion protein with an AD- or BD-tag either at the N- or C-terminus. All constructed plasmids are listed in Table S2. Transformation of yeast cells was performed according to manufacturer’s guidelines using the Frozen-EZ Yeast Transformation II Kit (Zymo Research) and cells were selected on complete supplement mixture (CSM) lacking leucine and tryptophan (-Leu -Trp) dropout medium (MP Biochemicals) at 30 °C for 3–4 days. Y190 (Clontech) cells were used for measuring β-galactosidase activity. Formed colonies were spotted on a second plate (CSM -Leu -Trp) and incubated for 2 days. Afterwards a colony-lift filter assay was performed as described by Breeden *et al*. (54). A detailed protocol can be found on protocols.io (https://dx.doi.org/10.17504/protocols.io.v7ve9n6).

### Bacterial strains and growth conditions

Wild type *Synechocystis* sp. PCC 6803 (PCC-M, re-sequenced, (46)) and the *kaiC3* deletion mutant (40) were cultured photoautotrophically in BG11 medium (55) supplemented with 10 mM TES buffer (pH 8) under constant illumination with 50 µmol photons m^−2^s^−1^ of white light (Philips TLD Super 80/840) at 30 °C. Cells were grown either in Erlenmeyer flasks with constant shaking (140 rpm) or on plates (0.75 % Bacto-Agar, Difco) supplemented with 0.3 % thiosulfate. Detailed recipes can be found on protocols.io (https://dx.doi.org/10.17504/protocols.io.wj5fcq6).

### Spot assays

Experiments were performed as previously described (56). Strains were propagated mixotrophically on BG11 agar plates with the addition of 0.2 % (w/v) glucose. Dilution series of cell cultures started with OD_750nm_ 0.2 and OD_750nm_ 0.4, followed by incubation of the plates for 6 or 28 days under constant light conditions and in complete darkness, respectively. Plates were scanned and single spot intensities were quantified using Quantity One (Bio-Rad). Measured intensities were normalized to the intensity of the wild-type spot of the respective control grown under continuous light.

## Acknowledgement

We thank Junko Moriwaki, Nancy Sauer, Jennifer Andres, Lukas Pohlig, Thomas Volkmer and Megumi Fujimoto for technical assistance. This work was funded by the Deutsche Forschungsgemeinschaft (DFG, German Research Foundation) under Germanýs Excellence Strategy – EXC 2048/1 – Project ID: 390686111 to A. Wiegard and I.M. Axmann. The work was financially supported by Heine Research Academies - Travel Grants, and EMBO ASTF 145 to A. Wiegard, by grants AX 84/1-3 and EXC 1028 from the German Research Foundation to I.M. Axmann, by JSPS Grants-in-Aid 16H00784, 17K19247 and 19K05833 to K. Terauchi, and by grant WI2014/5-3 and WI2014/10-1 from the German Research Foundation to A. Wilde.

## Author contributions

I.M. Axmann, A. Wilde, A. Wiegard, C. Köbler and K. Terauchi conceived the project. A. Wiegard, C. Köbler A.K. Dörrich and K. Oyama designed, performed and analyzed experiments. I.M Axmann, A. Wilde and K. Terauchi supervised the study. All authors interpreted and discussed the data. A. Wiegard, C. Köbler and A. Wilde wrote the manuscript. I.M. Axmann, C. Azai, K. Terauchi, A.K. Dörrich, and K. Oyama commented essentially on the manuscript. All authors approved the manuscript.

## Supplementary methods

### Pull down analysis of KaiB and KaiC proteins

GST-KaiB1 and GST-KaiB3 proteins were expressed in *E. coli* BL21 using the pGEX-6P1 vector and extracted as described in the main text. Protein concentration of the generated whole cell lysate was determined using the Bradford method (Bradford MM. 1976. Anal Biochem 72:248-54). For immobilization of each KaiB protein, a volume corresponding to 40 mg whole protein content was incubated with 100 µl glutathione sepharose 4B (GE Healthcare) for 20 min at room temperature. The resin was thoroughly washed four times with extraction buffer [50 mM Tris/HCl (pH8), 150 mM NaCl, 0.5 mM EDTA, 1 mM DTT (+ 5 mM MgCl_2_, 1 mM ATP for KaiC proteins)] and subsequently incubated with *Synechocystis* wild type whole cell lysate, which had been generated from 200 ml cell culture grown to an OD_750nm_ of ~0.8. Briefly, cells were lysed in cold thylakoid buffer [50 mM Hepes/NaOH (pH7), 5 mM MgCl_2_, 25 mM CaCl_2_, 10% Glycerol] using glass beads (mixture of 0.1–0.11 mm and 0.25-0.5 mm size) in a bead beater (Reetsch) at 4 °C. 1 ml of the lysate was used for incubation with glutathione sepharose-bound GST-KaiB1 and GST-KaiB3 proteins to co-precipitate Kai interaction partners. Afterwards, the resin material was thoroughly washed with extraction buffer, mixed with SDS loading dye and incubated at 50 °C for 30 min before being subjected to SDS-PAGE and Western Blot analysis.

Expression and purification of 3xFLAG-tagged KaiC1 and KaiC3 proteins was performed as described earlier (Wiegard A, Dörrich AK, Deinzer HT, Beck C, Wilde A, Holtzendorff J, Axmann IM. 2013. Microbiology 159:948-958) from *Synechocystis*. After binding of 3xFLAG-KaiC1/3 to ANTI- FLAG M2 affinity gel (Sigma Aldrich) over night at 4 °C, the resin was thoroughly washed with cold FLAG buffer containing 0.03 % β-DM, followed by one washing step without the addition of β-DM. For pull down of KaiB proteins, 150 µl resin-bound 3x-FLAG-KaiC1 and 3xFLAG-KaiC3 were incubated with 100 µl of *E. coli* BL21 whole cell lysate (adjusted to a protein content of 10 µg/µl) containing GST-KaiB1 and GST-KaiB3, respectively. After removing unbound proteins by washing with FLAG buffer, the samples were subjected to SDS-PAGE and Western Blot analysis.

### Antibodies

Antibodies against KaiB1 and KaiB3 were produced as whole-rabbit-IgG fraction by Pineda Antikörper Services using synthetic peptides coupled to keyhole limpet haemocyanin as antigens. For the synthetic peptides, epitopes were completed N- and C-terminally to 15 AA. The peptide sequences were CIDVLK**NPQLAEEDKIL**AT (bold epitope corresponds to AA 50-59 in KaiB1) and CLDIVPEG**LQVRL**PED (bold epitope corresponds to AA 95-98 in KaiB3), respectively. For detection of KaiC1 and KaiC3 the specific peptide derived antibodies described in Wiegard *et al*. were used (Wiegard A, Dörrich AK, Deinzer HT, Beck C, Wilde A, Holtzendorff J, Axmann IM. 2013. Microbiology 159:948-958). HemA antibody was produced by Pineda Antikörper Services using the recombinant protein and has been described in Sobotka *et al*. (Sobotka R, Tichy M, Wilde A, Hunter CN. 2011. Plant Physiol 155:1735-47). The antibody against AtpB was kindly provided by K.-D. Irrgang (Technical University Berlin, Germany).

---

**Table S1.**
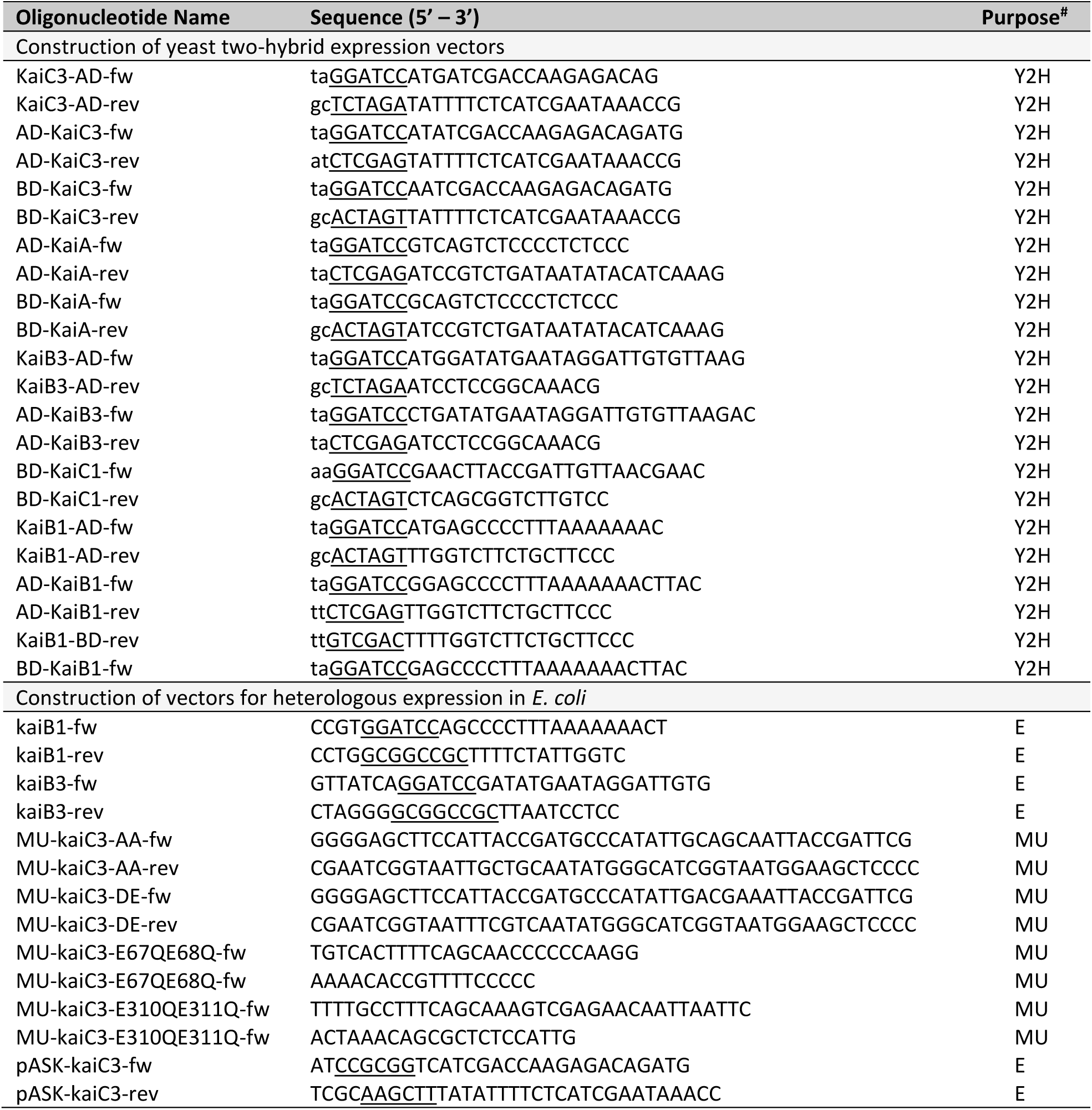
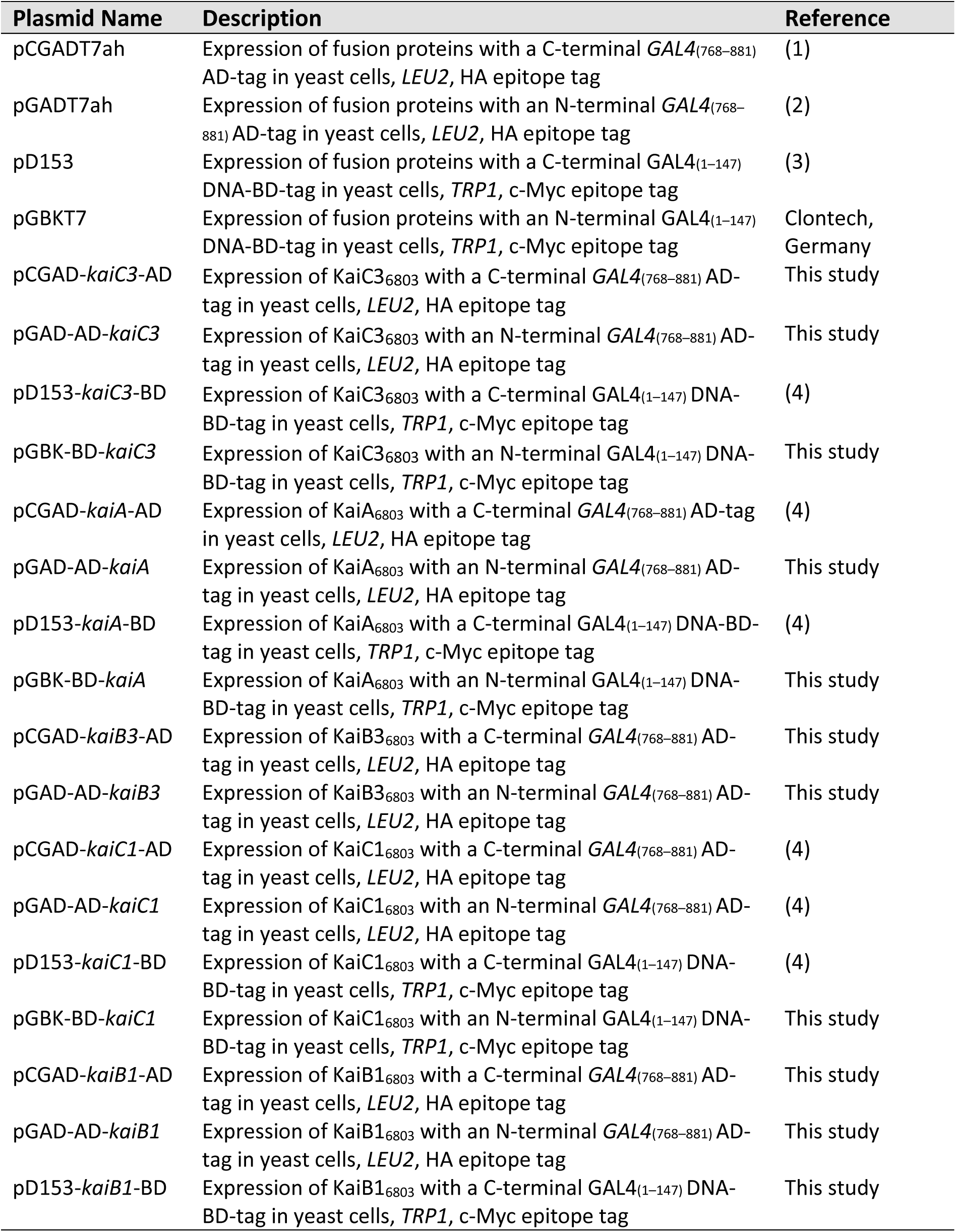
Oligonucleotides used in this study. Restriction sites are underlined. ^#^ Y2H, expression in yeast cells; E, expression in *E. coli* cells; MU, mutagenesis

**Table S2.**
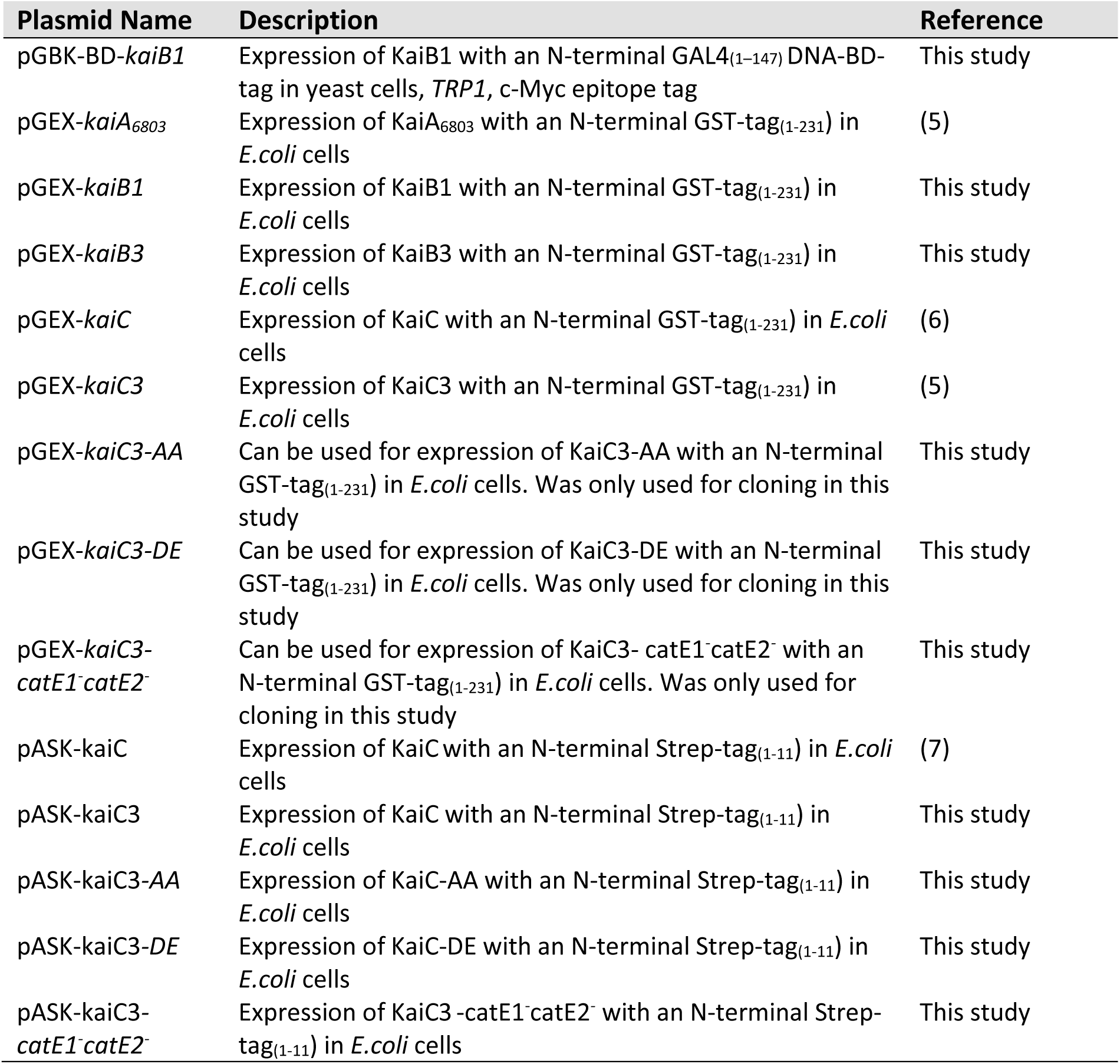
Plasmids used in this study.

**Table S2 references**

1. Rausenberger J, Tscheuschler A, Nordmeier W, Wust F, Timmer J, Schafer E, Fleck C, Hiltbrunner A. 2011. Photoconversion and nuclear trafficking cycles determine phytochrome A’s response profile to far-red light. Cell 146:813-25.

2. Hiltbrunner A, Viczian A, Bury E, Tscheuschler A, Kircher S, Toth R, Honsberger A, Nagy F, Fankhauser C, Schafer E. 2005. Nuclear accumulation of the phytochrome A photoreceptor requires FHY1. Curr Biol 15:2125-30.

3. Shimizu-Sato S, Huq E, Tepperman JM, Quail PH. 2002. A light-switchable gene promoter system. Nat Biotechnol 20:1041-4.

4. Axmann IM, Dühring U, Seeliger L, Arnold A, Vanselow JT, Kramer A, Wilde A. 2009. Biochemical evidence for a timing mechanism in Prochlorococcus. J Bacteriol 191:5342-7.

5. Wiegard A, Dörrich AK, Deinzer HT, Beck C, Wilde A, Holtzendorff J, Axmann IM. 2013. Biochemical analysis of three putative KaiC clock proteins from Synechocystis sp. PCC 6803 suggests their functional divergence. Microbiology 159:948-958.

6. Nishiwaki T, Satomi Y, Nakajima M, Lee C, Kiyohara R, Kageyama H, Kitayama Y, Temamoto M, Yamaguchi A, Hijikata A, Go M, Iwasaki H, Takao T, Kondo T. 2004. Role of KaiC phosphorylation in the circadian clock system of Synechococcus elongatus PCC 7942. Proc Natl Acad Sci U S A 101:13927-32.

7. Oyama K, Azai C, Nakamura K, Tanaka S, Terauchi K. 2016. Conversion between two conformational states of KaiC is induced by ATP hydrolysis as a trigger for cyanobacterial circadian oscillation. Sci Rep 6:32443.

**Figure S1.**
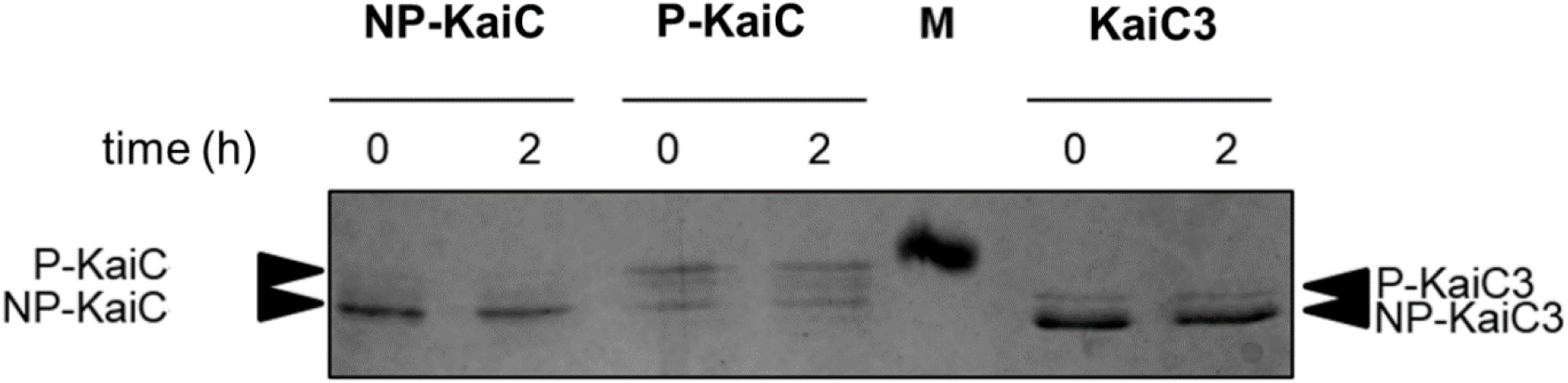
Phosphorylation level of KaiC proteins used for ATP synthase activity assay shown in Fig. 1. Proteins were separated via SDS-PAGE using a polyacrylamide gel with 11 %T, 0.67 %C (see dx.doi.org/10.17504/protocols.io.gysbxwe for method description). Fully phosphorylated (P-KaiC) and dephosphorylated (NP-KaiC) *S. elongatus* KaiC proteins were generated by incubating the protein for 2 weeks at 4 °C or overnight at 30 °C, respectively.

**Figure S2.**
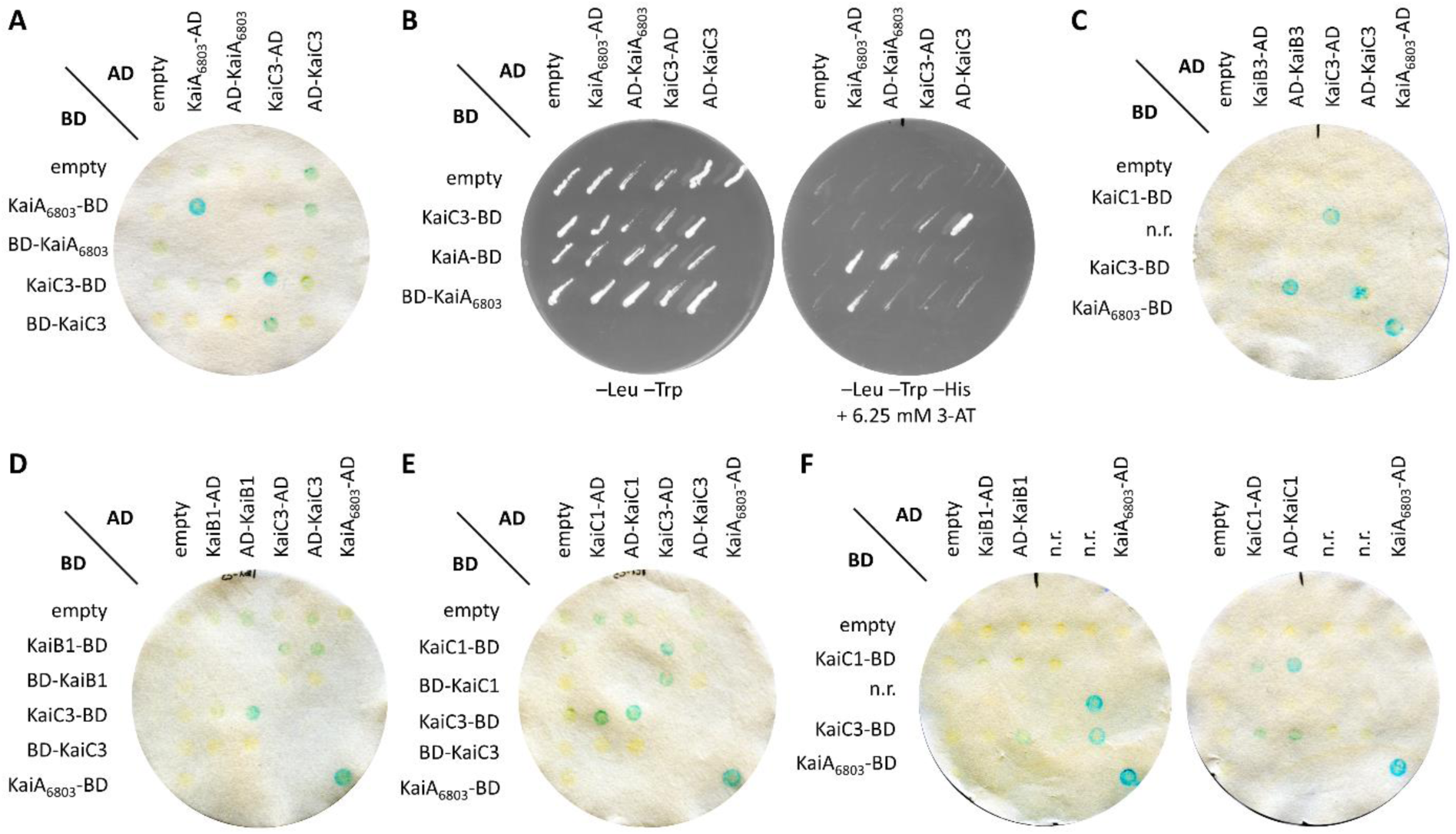
Original scans of the KaiC3 interaction with KaiB3 and the proteins of the main oscillator KaiC1, KaiB1. Yeast two-hybrid reporter strains carrying the respective bait and prey plasmids, were selected by plating on complete supplement medium (CSM) lacking leucine and tryptophan (-Leu -Trp). As a positive control, KaiA_6803_ dimer interaction was used. AD, GAL4 activation domain; BD, GAL4 DNA-binding domain; n.r., interactions not relevant for this study. **A, C-F**: Physical interaction between bait and prey fusion proteins is indicated by a color change in the assays using 5-brom-4-chlor-3-indoxyl-β-D-galactopyranoside. **B:** Physical interaction between bait and prey fusion proteins is determined by growth on complete medium lacking leucine, tryptophan and histidine (-Leu -Trp -His) and addition of 6.25 mM 3-amino-1,2,4-triazole (3-AT). A detailed protocol can be found on protocols.io (https://dx.doi.org/10.17504/protocols.io.wcnfave).

**Figure S3.**
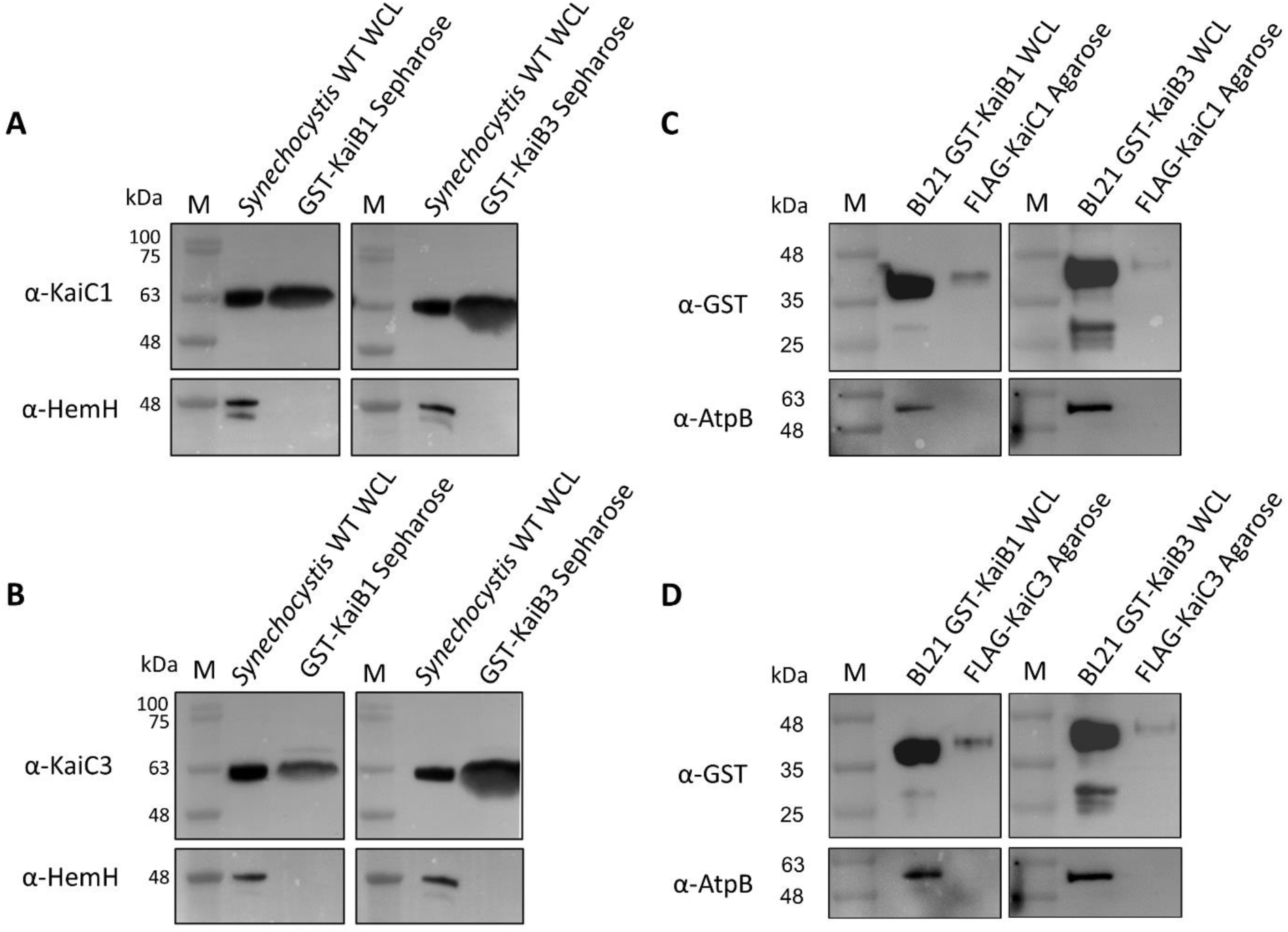
Interaction of *Synechocystis* KaiB and KaiC proteins in pull down analysis. **A,B** GST-tagged KaiB1 and KaiB3 proteins were expressed in *E. coli* BL21 cells, bound to glutathione sepharose and incubated with *Synechocystis* WT whole cell lysate (WCL). In the eluate, KaiC1 (A) and KaiC3 (B) were detected by Western Blot analysis using specific antibodies. As negative control, blots were incubated with an antiserum against the ferrochelatase HemH, because we did not expect an interaction between the KaiB proteins and HemH. **C,D** FLAG- KaiC1 (C) and FLAG- KaiC3 (D) were expressed in *Synechocystis*, bound to Anti-FLAG-agarose and incubated with whole cell lysate (WCL) from *E. coli* BL21 cells expressing GST-KaiB1 and GST-KaiB3 proteins, respectively. To detect whether GST-KaiB proteins were co-eluted we used an antibody raised against GST. Incubation with an AtpB antibody served as negative control. Both experiments were performed once.

**Figure S4.**
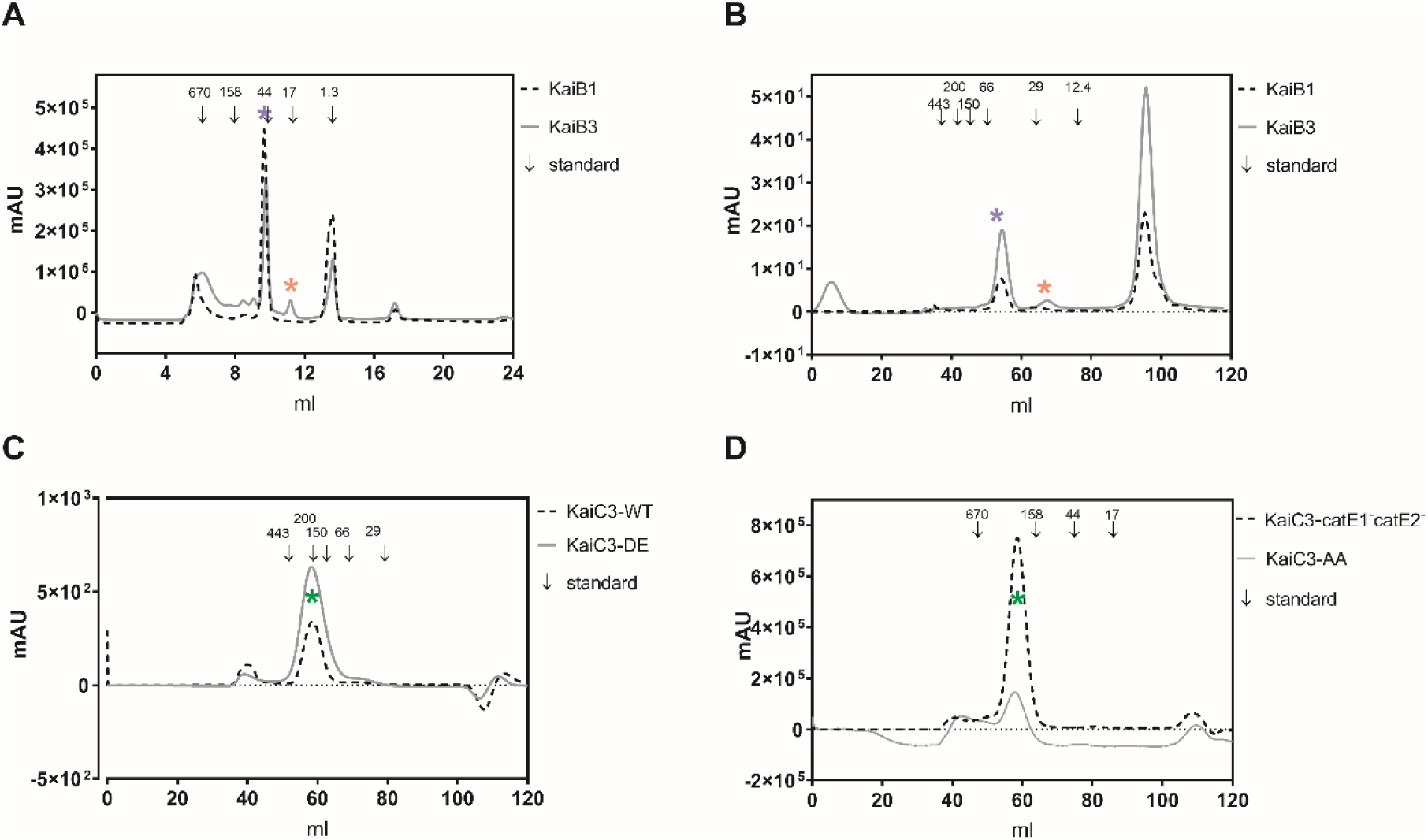
Purification of KaiB and KaiC3 proteins used in this study. **AB:** KaiB proteins were subsequently purified via affinity chromatography, anion exchange chromatography and size exclusion chromatography. Shown are the chromatograms after size exclusion chromatography using a superdex 200 Increase 10/30 GL column (A) or Sephacryl S200 HR HiPrep 16/60 column (B). On both columns KaiB3 was separated into a monomer (red asterisk) and tetramer (blue asterisk), whereas KaiB1 was mainly eluted as tetramer (blue asterisk). **CD:** KaiC3 proteins were purified via affinity chromatography followed by size exclusion chromatography on a Sephacryl S300 HR HiPrep 16/60 Sephacryl column. All KaiC3 proteins eluted as oligomer (green asterisk). Arrows indicate the size of standard proteins in kDa.

**Figure S5.**
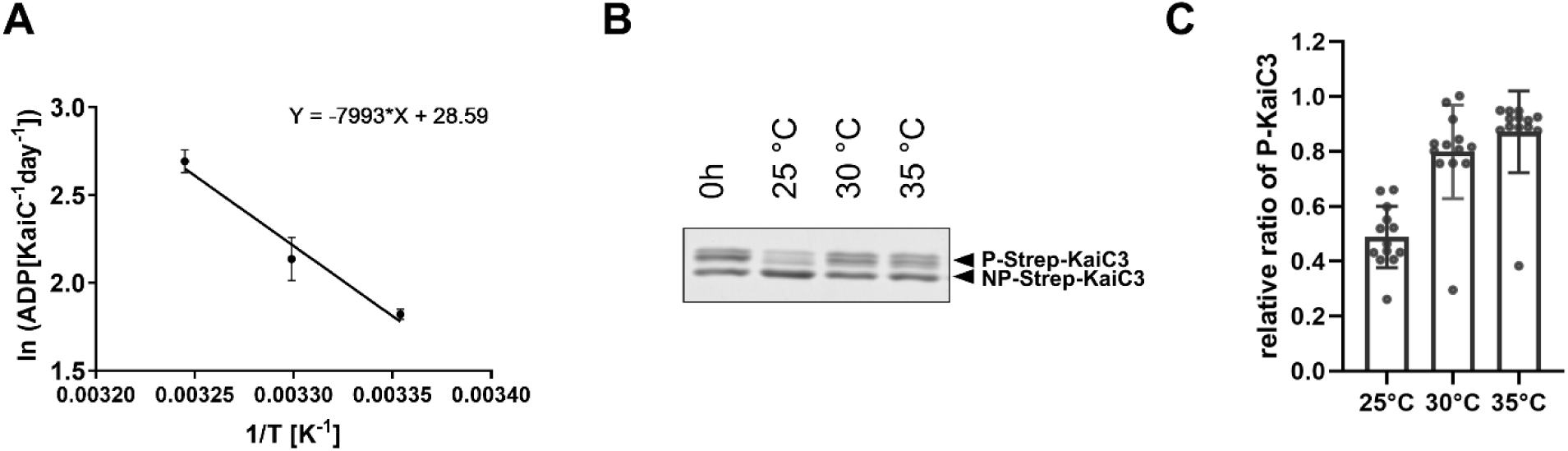
ATPase activity and dephosphorylation of KaiC3 are temperature dependent. **A.** Temperature-dependent ATPase activity of Strep-KaiC3 (also displayed in Fig. 4B) shown in an Arrhenius plot. Strep-KaiC3 was incubated for 24 hours at 25°C, 30°C and 35°C and ADP production per day and monomer Strep-KaiC3 was calculated. To determine the activation energy, the natural logarithm of the mean ADP production rate was plotted against the reciprocal temperature. The activation energy of KaiC3 ATPase was calculated from the slope of a linear regression equation as E_a_=66.5 x kJ mol^−1^. B,C. Relative dephosphorylation of Strep-KaiC3. Strep-KaiC3 was incubated for 24 hours in 20 mM Tris-HCl/pH8, 150 mM NaCl, 5 mM MgCl2, 1 mM ATP at the indicated temperatures. Proteins were separated via SDS-PAGE using a polyacrylamide gel with 11 %T, 0.67 %C. The phosphorylation level at each temperature was determined as the ratio of P-Strep-KaiC3 to total Strep-KaiC3 (P-Strep-KaiC3 + NP-Strep-KaiC3) using ImageJ. A representative gel image is shown in **B**. In **C**, the relative phosphorylation levels compared to the stock protein (0h) of 5 experiments are plotted as mean values with standard deviation. Incubation at 25°C resulted in lower phosphorylation levels, demonstrating higher net-dephosphorylation.

## References

1. Wiegard A, Köbler C, Oyama, K, Dörrich, AK, Azai, C, Terauchi, K, Wilde, A, Axmann, IM. 2019. Synechocystis KaiC3 displays temperature and KaiB dependent ATPase activity and is important for viability in darkness. bioRxiv doi:10.1101/700500:700500.

2. Ditty JL, Williams SB, Golden SS. 2003. A cyanobacterial circadian timing mechanism. Annu Rev Genet 37:513–43.

3. Nakajima M, Imai K, Ito H, Nishiwaki T, Murayama Y, Iwasaki H, Oyama T, Kondo T. 2005. Reconstitution of circadian oscillation of cyanobacterial KaiC phosphorylation *in vitro*. Science 308:414–5.

4. Iwasaki H, Nishiwaki T, Kitayama Y, Nakajima M, Kondo T. 2002. KaiA-stimulated KaiC phosphorylation in circadian timing loops in cyanobacteria. Proc Natl Acad Sci U S A 99:15788–93.

5. Terauchi K, Kitayama Y, Nishiwaki T, Miwa K, Murayama Y, Oyama T, Kondo T. 2007. ATPase activity of KaiC determines the basic timing for circadian clock of cyanobacteria. Proc Natl Acad Sci U S A 104:16377–81.

6. Kitayama Y, Iwasaki H, Nishiwaki T, Kondo T. 2003. KaiB functions as an attenuator of KaiC phosphorylation in the cyanobacterial circadian clock system. EMBO J 22:2127–34.

7. Hayashi F, Suzuki H, Iwase R, Uzumaki T, Miyake A, Shen JR, Imada K, Furukawa Y, Yonekura K, Namba K, Ishiura M. 2003. ATP-induced hexameric ring structure of the cyanobacterial circadian clock protein KaiC. Genes Cells 8:287–96.

8. Mori T, Saveliev SV, Xu Y, Stafford WF, Cox MM, Inman RB, Johnson CH. 2002. Circadian clock protein KaiC forms ATP-dependent hexameric rings and binds DNA. Proc Natl Acad Sci U S A 99:17203–8.

9. Pattanayek R, Wang J, Mori T, Xu Y, Johnson CH, Egli M. 2004. Visualizing a circadian clock protein: crystal structure of KaiC and functional insights. Mol Cell 15:375–88.

10. Nishiwaki T, Iwasaki H, Ishiura M, Kondo T. 2000. Nucleotide binding and autophosphorylation of the clock protein KaiC as a circadian timing process of cyanobacteria. Proc Natl Acad Sci U S A 97:495–9.

11. Murakami R, Miyake A, Iwase R, Hayashi F, Uzumaki T, Ishiura M. 2008. ATPase activity and its temperature compensation of the cyanobacterial clock protein KaiC. Genes Cells 13:387–95.

12. Nishiwaki T, Kondo T. 2012. Circadian autodephosphorylation of cyanobacterial clock protein KaiC occurs via formation of ATP as intermediate. J Biol Chem 287:18030–5.

13. Egli M, Mori T, Pattanayek R, Xu Y, Qin X, Johnson CH. 2012. Dephosphorylation of the core clock protein KaiC in the cyanobacterial KaiABC circadian oscillator proceeds via an ATP synthase mechanism. Biochemistry 51:1547–58.

14. Phong C, Markson JS, Wilhoite CM, Rust MJ. 2013. Robust and tunable circadian rhythms from differentially sensitive catalytic domains. Proc Natl Acad Sci U S A 110:1124–9.

15. Oyama K, Azai C, Nakamura K, Tanaka S, Terauchi K. 2016. Conversion between two conformational states of KaiC is induced by ATP hydrolysis as a trigger for cyanobacterial circadian oscillation. Sci Rep 6:32443.

16. Shultzaberger RK, Boyd JS, Diamond S, Greenspan RJ, Golden SS. 2015. Giving Time Purpose: The Synechococcus elongatus Clock in a Broader Network Context. Annu Rev Genet 49:485–505.

17. Dong G, Yang Q, Wang Q, Kim YI, Wood TL, Osteryoung KW, van Oudenaarden A, Golden SS. 2010. Elevated ATPase activity of KaiC applies a circadian checkpoint on cell division in *Synechococcus elongatus*. Cell 140:529–39.

18. Swan JA, Golden SS, LiWang A, Partch CL. 2018. Structure, function, and mechanism of the core circadian clock in cyanobacteria. J Biol Chem 293:5026–5034.

19. Diamond S, Jun D, Rubin BE, Golden SS. 2015. The circadian oscillator in Synechococcus elongatus controls metabolite partitioning during diurnal growth. Proc Natl Acad Sci U S A 112:E1916–25.

20. Welkie DG, Rubin BE, Diamond S, Hood RD, Savage DF, Golden SS. 2019. A Hard Day’s Night: Cyanobacteria in Diel Cycles. Trends Microbiol 27:231–242.

21. Pattanayak G, Rust MJ. 2014. The cyanobacterial clock and metabolism. Curr Opin Microbiol 18:90–5.

22. Woelfle MA, Ouyang Y, Phanvijhitsiri K, Johnson CH. 2004. The adaptive value of circadian clocks: an experimental assessment in cyanobacteria. Curr Biol 14:1481–6.

23. Köbler C, Schultz SJ, Kopp D, Voigt K, Wilde A. 2018. The role of the Synechocystis sp. PCC 6803 homolog of the circadian clock output regulator RpaA in day-night transitions. Mol Microbiol 110:847–861.

24. Ungerer J, Wendt KE, Hendry JI, Maranas CD, Pakrasi HB. 2018. Comparative genomics reveals the molecular determinants of rapid growth of the cyanobacterium Synechococcus elongatus UTEX 2973. Proc Natl Acad Sci U S A doi:10.1073/pnas.1814912115.

25. Welkie DG, Rubin BE, Chang YG, Diamond S, Rifkin SA, LiWang A, Golden SS. 2018. Genome-wide fitness assessment during diurnal growth reveals an expanded role of the cyanobacterial circadian clock protein KaiA. Proc Natl Acad Sci U S A 115:E7174–e7183.

26. Diamond S, Rubin BE, Shultzaberger RK, Chen Y, Barber CD, Golden SS. 2017. Redox crisis underlies conditional light-dark lethality in cyanobacterial mutants that lack the circadian regulator, RpaA. Proc Natl Acad Sci U S A 114:E580–e589.

27. Shih PM, Wu D, Latifi A, Axen SD, Fewer DP, Talla E, Calteau A, Cai F, Tandeau de Marsac N, Rippka R, Herdman M, Sivonen K, Coursin T, Laurent T, Goodwin L, Nolan M, Davenport KW, Han CS, Rubin EM, Eisen JA, Woyke T, Gugger M, Kerfeld CA. 2013. Improving the coverage of the cyanobacterial phylum using diversity-driven genome sequencing. Proc Natl Acad Sci U S A 110:1053–8.

28. Schmelling NM, Lehmann R, Chaudhury P, Beck C, Albers S-VV, Axmann IMM, Wiegard A. 2017. Minimal Tool Set for a Prokaryotic Circadian Clock. bioRxiv doi:10.1101/075291.

29. Axmann IM, Hertel S, Wiegard A, Dorrich AK, Wilde A. 2014. Diversity of KaiC-based timing systems in marine Cyanobacteria. Mar Genomics 14C:3–16.

30. Aoki S, Onai K. 2009. Circadian clocks of Synechocystis sp. strain PCC 6803, Thermosynechococcus elongatus, Prochlorococcus spp., Trichodesmium spp. and other species, p 259–282. *In* Ditty JL, Mackey SR, Johnson CH (ed), Bacterial circadian programs. Springer Berlin Heidelberg.

31. Dvornyk V, Vinogradova O, Nevo E. 2003. Origin and evolution of circadian clock genes in prokaryotes. Proc Natl Acad Sci U S A 100:2495–500.

32. Wiegard A, Dörrich AK, Deinzer HT, Beck C, Wilde A, Holtzendorff J, Axmann IM. 2013. Biochemical analysis of three putative KaiC clock proteins from Synechocystis sp. PCC 6803 suggests their functional divergence. Microbiology 159:948–958.

33. Johnson CH, Egli M. 2014. Metabolic compensation and circadian resilience in prokaryotic cyanobacteria. Annu Rev Biochem 83:221–47.

34. Holtzendorff J, Partensky F, Mella D, Lennon JF, Hess WR, Garczarek L. 2008. Genome streamlining results in loss of robustness of the circadian clock in the marine cyanobacterium *Prochlorococcus marinus* PCC 9511. J Biol Rhythms 23:187–99.

35. Kanesaki Y, Shiwa Y, Tajima N, Suzuki M, Watanabe S, Sato N, Ikeuchi M, Yoshikawa H. 2012. Identification of substrain-specific mutations by massively parallel whole-genome resequencing of Synechocystis sp. PCC 6803. DNA Res 19:67–79.

36. Dvornyk V, Knudsen B. 2005. Functional divergence of the circadian clock proteins in prokaryotes. Genetica 124:247–54.

37. Axmann IM, Dühring U, Seeliger L, Arnold A, Vanselow JT, Kramer A, Wilde A. 2009. Biochemical evidence for a timing mechanism in *Prochlorococcus*. J Bacteriol 191:5342–7.

38. Ma P, Mori T, Zhao C, Thiel T, Johnson CH. 2016. Evolution of KaiC-Dependent Timekeepers: A Proto-circadian Timing Mechanism Confers Adaptive Fitness in the Purple Bacterium Rhodopseudomonas palustris. PLoS Genet 12:e1005922.

39. Mukaiyama A, Ouyang D, Furuike Y, Akiyama S. 2019. KaiC from a cyanobacterium Gloeocapsa sp. PCC 7428 retains functional and structural properties required as the core of circadian clock system. International Journal of Biological Macromolecules 131:67–73.

40. Dörrich AK, Mitschke J, Siadat O, Wilde A. 2014. Deletion of the Synechocystis sp. PCC 6803 kaiAB1C1 gene cluster causes impaired cell growth under light-dark conditions. Microbiology doi:10.1099/mic.0.081695-0.

41. Kucho K-i, Okamoto K, Tsuchiya Y, Nomura S, Nango M, Kanehisa M, Ishiura M. 2005. Global analysis of circadian expression in the cyanobacterium Synechocystis sp. strain PCC 6803. J Bacteriol 187:2190–9.

42. Beck C, Hertel S, Rediger A, Lehmann R, Wiegard A, Kolsch A, Heilmann B, Georg J, Hess WR, Axmann IM. 2014. A daily expression pattern of protein-coding genes and small non-coding RNAs in Synechocystis sp. PCC 6803. Appl Environ Microbiol doi:10.1128/AEM.01086-14.

43. van Alphen P, Hellingwerf KJ. 2015. Sustained Circadian Rhythms in Continuous Light in Synechocystis sp. PCC6803 Growing in a Well-Controlled Photobioreactor. PLoS One 10:e0127715.

44. Saha R, Liu D, Hoynes-O’Connor A, Liberton M, Yu J, Bhattacharyya-Pakrasi M, Balassy A, Zhang F, Moon TS, Maranas CD, Pakrasi HB. 2016. Diurnal Regulation of Cellular Processes in the Cyanobacterium Synechocystis sp. Strain PCC 6803: Insights from Transcriptomic, Fluxomic, and Physiological Analyses. MBio 7.

45. Zavřel T, Očenášová P, Červený J. 2017. Phenotypic characterization of Synechocystis sp. PCC 6803 substrains reveals differences in sensitivity to abiotic stress. PLoS One 12:e0189130–e0189130.

46. Trautmann D, Voss B, Wilde A, Al-Babili S, Hess WR. 2012. Microevolution in cyanobacteria: re-sequencing a motile substrain of Synechocystis sp. PCC 6803. DNA Res 19:435–48.

47. Anderson SL, McIntosh L. 1991. Light-activated heterotrophic growth of the cyanobacterium Synechocystis sp. strain PCC 6803: a blue-light-requiring process. J Bacteriol 173:2761–7.

48. Loza-Correa M, Sahr T, Rolando M, Daniels C, Petit P, Skarina T, Gomez Valero L, Dervins-Ravault D, Honore N, Savchenko A, Buchrieser C. 2014. The Legionella pneumophila kai operon is implicated in stress response and confers fitness in competitive environments. Environ Microbiol 16:359–81.

49. Snijder J, Schuller JM, Wiegard A, Lossl P, Schmelling N, Axmann IM, Plitzko JM, Forster F, Heck AJ. 2017. Structures of the cyanobacterial circadian oscillator frozen in a fully assembled state. Science 355:1181–1184.

50. Tseng R, Goularte NF, Chavan A, Luu J, Cohen SE, Chang YG, Heisler J, Li S, Michael AK, Tripathi S, Golden SS, LiWang A, Partch CL. 2017. Structural basis of the day-night transition in a bacterial circadian clock. Science 355:1174–1180.

51. Chang YG, Cohen SE, Phong C, Myers WK, Kim YI, Tseng R, Lin J, Zhang L, Boyd JS, Lee Y, Kang S, Lee D, Li S, Britt RD, Rust MJ, Golden SS, LiWang A. 2015. Circadian rhythms. A protein fold switch joins the circadian oscillator to clock output in cyanobacteria. Science 349:324–8.

52. Hitomi K, Oyama T, Han S, Arvai AS, Getzoff ED. 2005. Tetrameric architecture of the circadian clock protein KaiB. A novel interface for intermolecular interactions and its impact on the circadian rhythm. J Biol Chem 280:19127–35.

53. Espinosa J, Boyd JS, Cantos R, Salinas P, Golden SS, Contreras A. 2015. Cross-talk and regulatory interactions between the essential response regulator RpaB and cyanobacterial circadian clock output. Proc Natl Acad Sci U S A 112:2198–203.

54. Breeden L, Nasmyth K. 1985. Regulation of the yeast HO gene. Cold Spring Harb Symp Quant Biol 50:643–50.

55. Rippka R, Deruelles J, Waterbury JB, Herdman M, Stanier RY. 1979. Generic assignments, strain histories and properties of pure cultures of cyanobacteria. J Gen Microbiol 111:1–61.

56. Dörrich AK, Wilde A. 2015. Spot Assays for Viability Analysis of Cyanobacteria. Bio-protocol 5:e1574.

